# Genetic background sets the trajectory of cancer evolution

**DOI:** 10.1101/2025.01.13.632787

**Authors:** Sarah J. Aitken, Frances Connor, Christine Feig, Tim F. Rayner, Margus Lukk, Juliet Luft, Stuart Aitken, Claudia Arnedo-Pac, James F. Hayes, Michael D. Nicholson, Vasavi Sundaram, Jan C. Verburg, John Connelly, Craig J. Anderson, Mikaela Behm, Susan Campbell, Maëlle Daunesse, Ailith Ewing, Vera B. Kaiser, Elissavet Kentepozidou, Oriol Pich, Aisling M. Redmond, Javier Santoyo-Lopez, Inés Sentís, Lana Talmane, Liver Cancer Evolution Consortium, Paul Flicek, Núria López-Bigas, Colin A. Semple, Martin S. Taylor, Duncan T. Odom

## Abstract

Human cancers are heterogeneous. Their genomes evolve from genetically diverse germlines in complex and dynamic environments, including exposure to potential carcinogens. This heterogeneity of humans, our environmental exposures, and subsequent tumours makes it challenging to understand the extent to which cancer evolution is predictable. Addressing this limitation, we re-ran early tumour evolution hundreds of times in diverse, inbred mouse strains, capturing genetic variation comparable to and beyond that found in human populations. The sex, environment, and carcinogenic exposures were all controlled and tumours comprehensively profiled with whole genome and transcriptome sequencing. Within a strain, there was a high degree of consistency in the mutational landscape, a limited range of driver mutations, and all strains converged on the acquisition of a MAPK activating mutation with similar transcriptional disruption of that pathway. Despite these similarities in the phenotypic state of tumours, different strains took markedly divergent paths to reach that state. This included pronounced biases in the precise driver mutations, the strain specific occurrence of whole genome duplication, and differences in subclonal selection that reflected both cancer susceptibility and tumour growth rate. These results show that interactions between the germline genome and the environment are highly deterministic for the trajectory of tumour genome evolution, and even modest genetic divergence can substantially alter selection pressures during cancer development, influencing both cancer risk and the biology of the tumour that develops.

## Main text

Among the most critical and yet least understood stages of cancer formation are the earliest. We lack a comprehensive account of how normal cells acquire driver mutations and interact with environmental factors to achieve oncogenic transformation^1–3^. Retrospective analysis of heterogeneous human tumours has inferred that a limited set of common drivers are involved in the initiation of most cancers^4^, and suggested that genetic background can influence cancer development^5,6^. Such studies are hampered by the diversity of the human population and poorly recorded histories of risk-factor exposure. As a consequence it is not clear how consistently cancer would develop given the same genetic background and same environment, or how germline differences influence the trajectory to tumourigenesis.

A powerful strategy to understand the mechanisms of tumorigenesis would be to implement Gould’s thought experiment of replaying evolution^7^, whereby cancer development is experimentally repeated hundreds of times for comprehensive analysis. Organoid based in vitro approaches^8^ provide an experimentally tractable system for early tumour development but they cannot capture the full in vivo context in which transformed cells are mutagenised and undergo selection. Here, we developed an in vivo prospective approach to study early cancer development, controlling for many of the genetic and environmental variables complicating human studies, and analysed how genetic background shapes both initial oncogenic transformation and subsequent gene-environment interactions.

Mutagenesis of mouse liver by the DNA damaging mutagen diethylnitrosamine (DEN) is a highly controlled system of *in vivo* tumorigenesis that reproducibly creates hepatocyte-derived tumours that histologically mimic human hepatocellular carcinoma (HCC)^9,10^. DEN is metabolically activated by centrilobular hepatocytes to form well-understood mutagenic DNA adducts^11^. Following a single DEN exposure 15 days after birth (P15), the C3H/HeOuJ mouse strain reliably develops multiple clonally distinct tumours within 25 weeks^12,13^, each tumour typically harbouring 60,000 base substitution mutations with characteristic DEN signatures. These tumours have activating mutations in one of the mitogen-activated protein kinase (MAPK) pathway genes *Hras*, *Braf*, *Egfr* or *Kras*, minimal structural variation, and do not have ongoing genome instability or mutator phenotypes^14^. A single discrete exposure to DEN results in a pronounced chromosome-scale mutational asymmetry through the process of lesion segregation, as tumours develop from the clonal growth of cells containing damage on only one strand of the DNA duplex^14^. The high density of strand-oriented mutations introduced in a single mutagen exposure is a powerful tool to analyse specific characteristics of tumourigenesis, including clonal dynamics, selection, homologous recombination events, and DNA repair dynamics^14–17^.

Here, using DEN we have re-run tumorigenesis hundreds of times in four inbred mouse strains and species with dramatic differences in cancer susceptibility^12,18–22^ (**Fig. 1a**). The natural genetic variation of these evolutionarily divergent inbred mice captures genetic variation equal to or exceeding that of human populations^23–25^ that has been historically neglected in population-level genetic studies^5,6^. By analysing the whole-genome sequences, paired transcriptomes, and histology of hundreds of DEN-induced tumours we demonstrate how cancer genome evolution is shaped by a cell’s starting genetic, epigenetic, and transcriptional states.

**Fig. 1.**
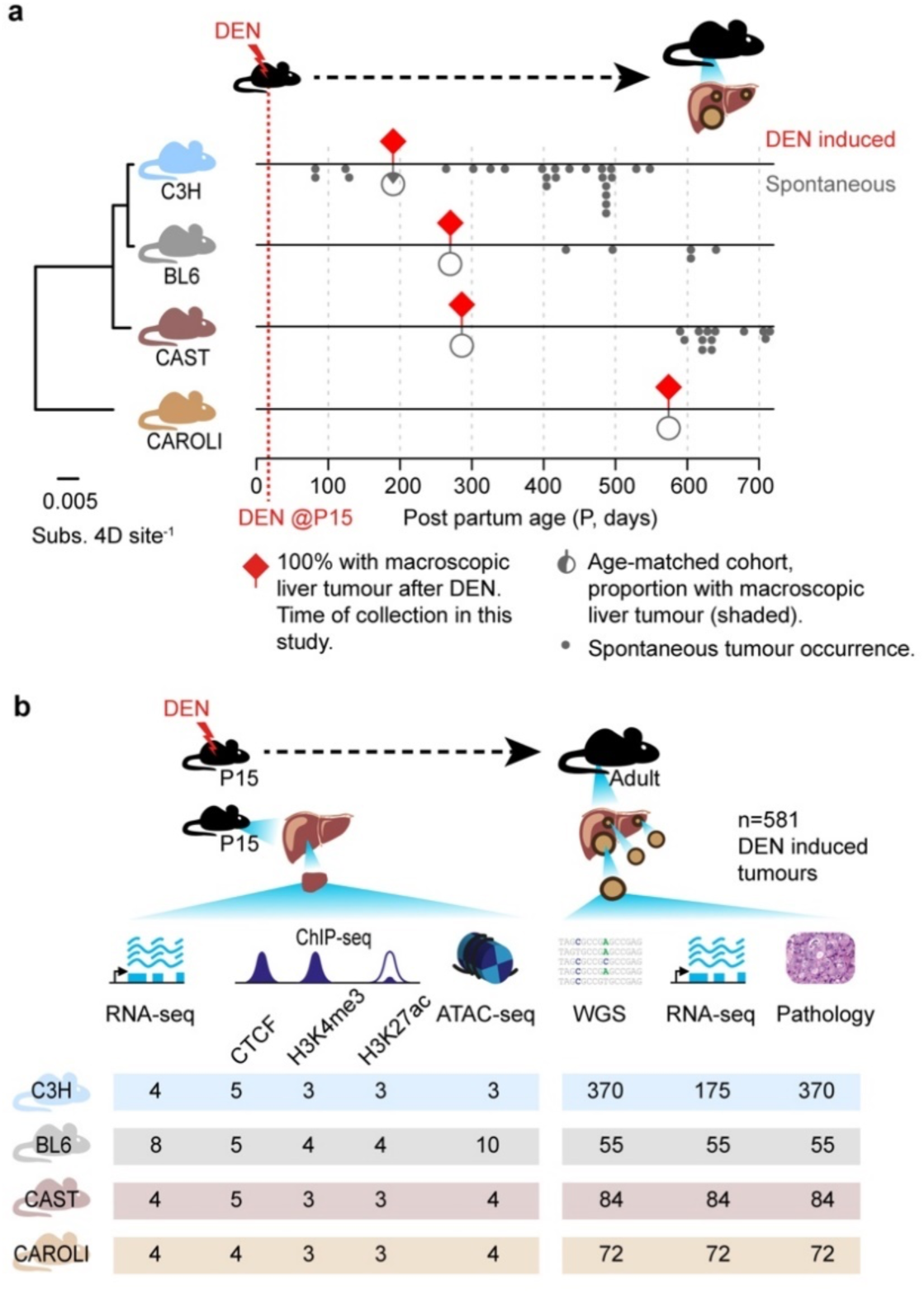
Cancer susceptibility is shaped by germline genetic variation. **a**, Summary of tumour induction by DEN and tumour latency in each mouse strain. Phylogenetic tree branch lengths are substitution rates of protein coding 4-fold degenerate sites. DEN-induced tumours were collected at fixed timepoints (x axis) for each strain (red diamonds), corresponding to 100% tumour incidence (Methods). Grey lollipops show the proportion (shaded area) of untreated, age-matched mice that developed spontaneous tumours at the same timepoint. Additional aged mice with spontaneous tumours are indicated as grey dots (**Supplementary Table 3**). **b**, Summary of study design and data generated. P15 mice were treated with DEN; control samples were collected from untreated mice for RNA-seq, ChIP-seq (10 individual livers pooled), and ATAC-seq. Liver tumours were isolated from adult mice at defined timepoints and subjected to WGS, RNA-seq, and histopathological analysis (**Supplementary Table 2**).

### Germline differences in tumour susceptibility

We chemically induced liver tumours^13,14^ by injecting a single intraperitoneal dose of diethylnitrosamine (DEN) on postnatal day 15 (P15) into inbred male laboratory mice of the following genetic backgrounds: *Mus musculus musculus* (C3H/HeOuJ, n=104 and C57BL/6J, n=12), *Mus musculus castaneus* (CAST/EiJ, n=54), and *Mus caroli* (n=45), hereafter referred to as C3H, BL6, CAST, and CAROLI, respectively (**Fig. 1a**). For simplicity, these strains, subspecies, and species are subsequently referred to as strains. The resultant tumours (n=581; **Supplementary Table 1**) were dissected, and processed in parallel for comprehensive profiling using whole genome sequencing (WGS), total transcriptome sequencing (RNA-seq), and histopathological analysis (**Supplementary Table 2**). Additional spontaneous liver tumours from a cohort of aged, untreated animals were collected and processed in parallel for comparison (**Supplementary Table 3**; n=39 of which underwent tumour profiling). Detailed characterisation of the transcriptional state and regulatory landscape of normal liver tissue from untreated P15 mice (the developmental time of DEN exposure) was performed using total RNA-seq, ATAC-seq, and ChIP-seq (**Fig. 1b**).

We measured a marked variation in susceptibility to DEN-induced tumourigenesis between strains (**Fig. 1a**). Following treatment, tumours were consistently present after 25 weeks (P190) in C3H but not until 36 weeks (P267) in BL6, in agreement with prior studies^12,19^. For the first time, we analysed the DEN model in CAST and CAROLI which revealed further extended latencies: 38 weeks (P281) in CAST, and not until 78 weeks (P561) in CAROLI. Spontaneous tumours in untreated mice follow the same trend in latency, with C3H most susceptible, then BL6, CAST, and CAROLI progressively more resistant (**Fig. 1a**; **Supplementary Table 3**). Collectively, these data constitute a unique, well powered, and highly controlled resource to study the molecular mechanisms underlying early tumourigenesis *in vivo*.

### Genetic background shapes genome stability

Given the striking effect of genetic background on tumour latency, we explored the associations of latency with underlying patterns of mutagenesis and genome stability by comparing the somatic changes in whole genome sequencing of DEN induced tumours between the four strains (total n=581). The type and sequence context of base substitution mutations (**Fig. 2a**) broadly correspond to the DEN1 and DEN2 mutation signatures derived from only C3H tumours^14^. However, mutation signature deconvolution involving all four strains reveals a phylogenetic shift in the DEN1 mutation signature, in which CAROLI has a decreased propensity for T→C mutations compared to the other strains (**Extended Data Fig. 1**).

**Fig. 2.**
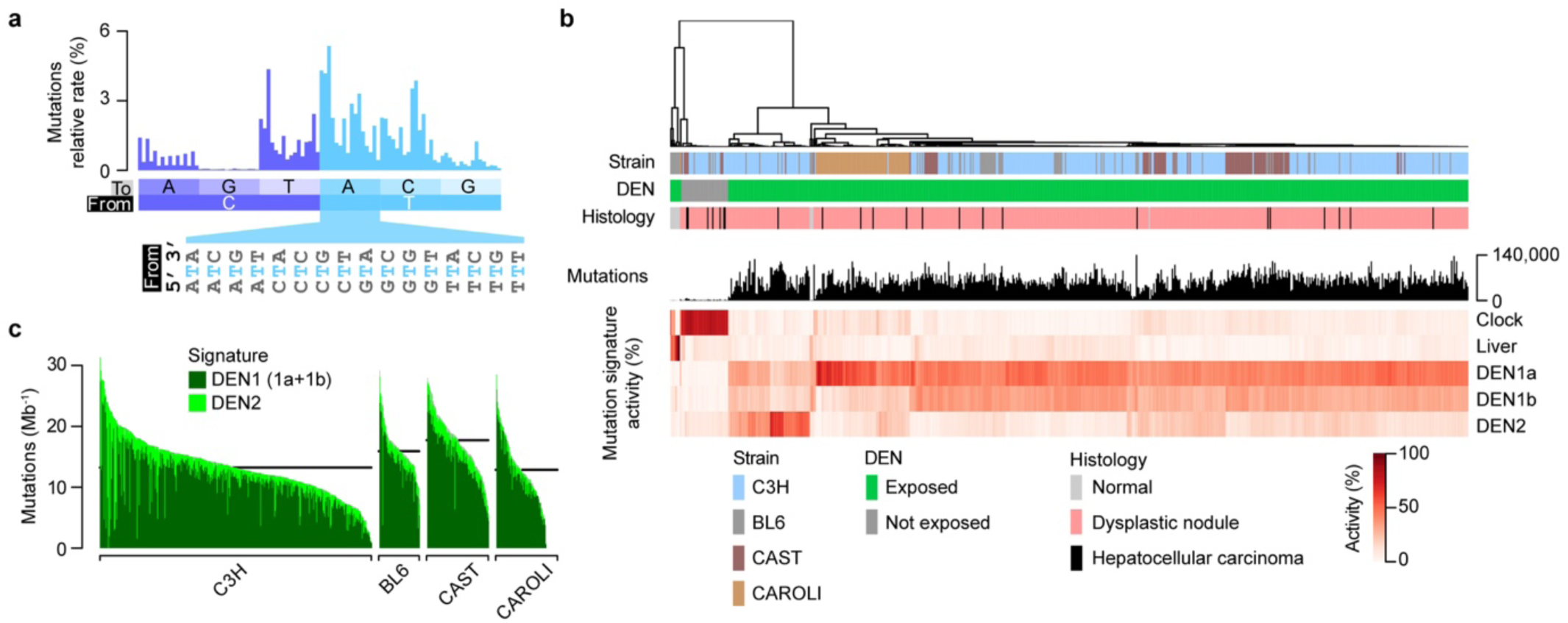
Genetic background subtly impacts the rate and spectrum of small mutations. **a**, Base substitution mutation rates calculated in aggregate over n=581 DEN induced tumours. Substitutions categorised by the type of change (from → to) conditioned on the identity of the upstream and downstream nucleotide, examples given for T→A mutations. Percentages included reverse complement mutations (e.g. T→C includes A→G). **b,** Mutation rate profiles (as panel **a**) were calculated for each whole genome sequenced sample and clustered by cosine distance (dendrogram). In addition to the DEN induced tumours, spontaneous tumours and histologically normal liver tissue samples were included for the comparison of signatures (total n=647, indicated in the DEN and histology tracks). Single base substitution mutation signatures were deconvolved (Methods) to define five component signatures. DEN induced tumours have a higher mutation load (black histogram) and distinct signature profile (heatmap) to non-DEN exposed tumours and DEN-exposed normal tissues. Amongst the DEN induced tumours, those from the CAROLI strain cluster together whereas the other strains are intermixed when clustered by mutation rate profile. **c**, Genome wide substitution mutation rate distribution for each strain. Despite greater tumour susceptibility, C3H does not have a systematically higher rate of mutations. The contribution of the DEN induced signatures is indicated. Horizontal line denotes the median mutation rate per strain.

Although DEN1 signatures dominate, the contribution of DEN2 varies between tumours and is more prevalent in C3H than the other strains (p=0.026, two-sided Mann-Whitney U test, C3H versus all other strains; **Fig. 2b**). This may be partly explained by strain differences in the expression of MGMT, an enzyme that can directly repair the O6-ethG adducts that are the likely source of DEN2 signature mutations^14^.

Genome-wide base substitution burden varies considerably within and between mouse strains (**Fig. 2b**), and is typically higher in CAST (median 17.6 Mb^−1^; 75,973 per tumour) and BL6 (16.6 Mb^−1^) than C3H (13.5 Mb^−1^) and CAROLI (13.3 Mb^−1^). Since mutation burden is discordant with tumour latency, the high susceptibility of C3H is not simply explained by greater accumulation of mutations.

Unlike abundant substitution mutations, small (<50bp) insertion and deletion (indel) mutations are rare in DEN induced tumours (1.4%, sd. 1.05 of the rate of substitutions). Typically, indels introduce less than 2 changes per tumour that alter the amino acid sequence (mean 1.77, sd. 0.99), whereas single-base substitutions introduce 400 to 800 such changes (**Fig. 2c**). Between tumours, the rate of indel mutations is positively correlated with nucleotide substitution rate (Pearson’s cor=0.21, p=3.9×10^−7^) and the ID83-style signature of indel mutations is highly conserved. However, there are strain preferences in the nucleotide composition of longer inserted and deleted sequences (**Extended Data Fig. 2**).

### Genetic background determines propensity to whole genome duplication

The vast majority of C3H (97%), BL6 (94%), and CAST (100%) tumours demonstrate the characteristic mutation asymmetry of lesion segregation^14^. In contrast, over a third (37%) of the CAROLI tumours appear mutationally symmetric (**Fig. 3a-c**). These symmetric CAROLI tumours are typified by a significantly higher mutation load than asymmetric tumours (mean 1.4 fold higher, p=0.0014, Mann-Whitney U test) and approximately half the cellularity (unadjusted for ploidy, mean 1.8 fold lower, p=5.8×10^−15^, Mann-Whitney U test). Collectively these observations are consistent with both daughter genomes of the originally mutagenised DNA duplex contributing equally to the sequenced tumour. In other words, the complementary mutational asymmetry of the two daughter genomes, arising from the damaged DNA duplex, systematically cancel out each other^15,17^.

**Fig. 3.**
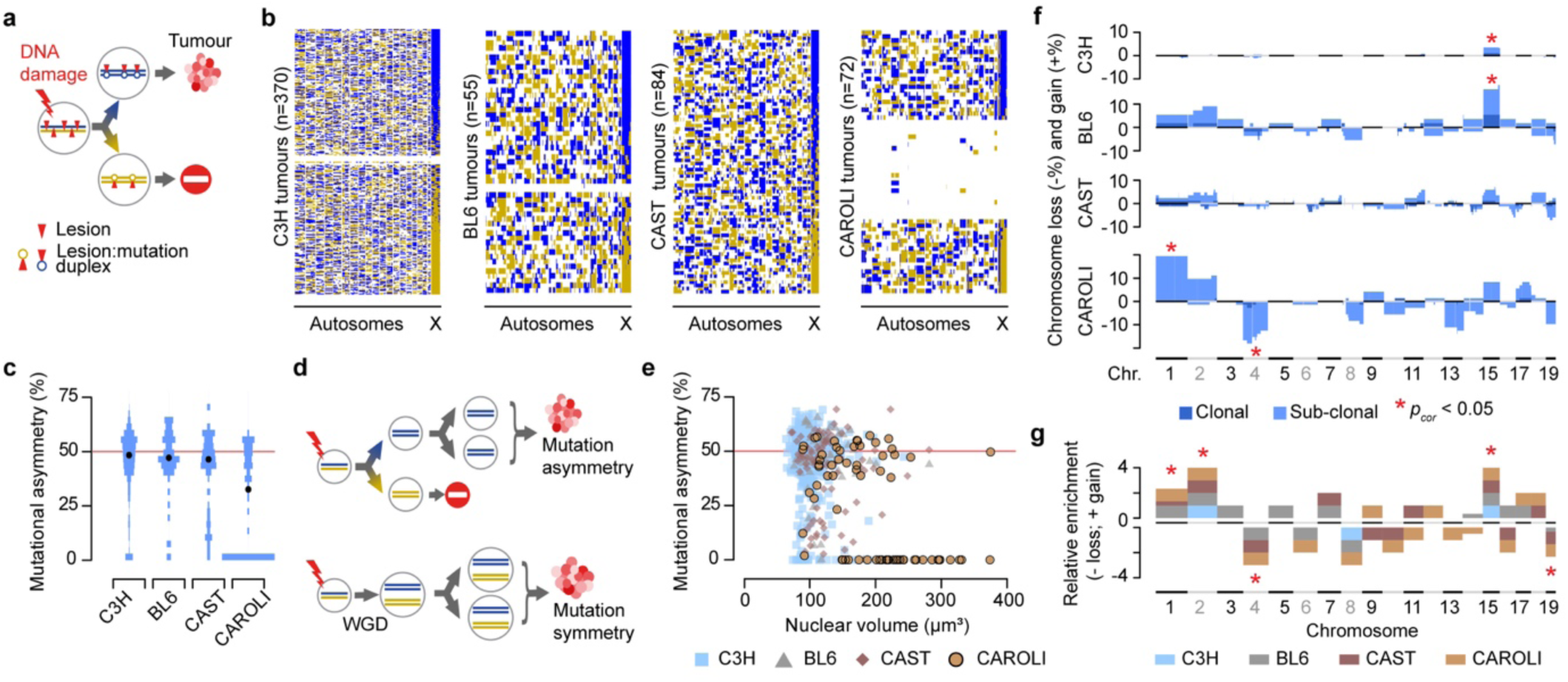
Profound strain differences in whole genome duplication during oncogenesis. **a**, Following DNA damaging exposure both strands of DNA are damaged but lesion segregation into daughter lineages and mutagenic replication over damage results in chromosome scale mutational asymmetry in the tumours that develop. **b**, Summary mutational asymmetry profiles of individual tumours (rows) across the genomes (x-axis) of each strain. Blue and gold represent mutationally asymmetric genomic segments (S>0.33, blue; S<-0.33 gold) and white represents no asymmetry. **c**, For strains other than CAROLI, the DEN induced tumours typically show mutational asymmetry across 50% of their autosomal genome (median, circle); in contrast, in CAROLI the distribution is bimodal, 36% with no or negligible asymmetry. **d**, The 50% asymmetry genomes are consistent with the classic lesion segregation model (upper schematic), whereas the mutationally symmetric genomes could be explained by the retention of both originally damaged genomes into the developed tumour, for example by whole genome duplication (WGD) in the first cell division following mutagenic insult (lower schematic). **e**, Approximate doubling of nuclear volume quantified from histology (x-axis) corresponds to the mutationally symmetric CAROLI tumours supporting a model of WGD. **f**, Large (>10 Mb) chromosomal gains (+%) and losses (−%) show pronounced differences in frequency between strains and significant directional enrichments for some chromosomes (* Bonferroni corrected two tailed permutation test p-value < 0.05). **g**, Whether a chromosome is preferentially gained or lost (y-axis relative enrichment metric, summed over strains) is conserved between strains. Three chromosomes are significantly enriched for copy-gains and two for copy loss in aggregate analysis over strains (* Bonferroni corrected two tailed permutation test p-value < 0.05).

One possible mechanistic explanation for these symmetric genomes is whole genome duplication (WGD) arising due to the failure of karyokinesis in the first mitotic division following mutagenesis, which would retain both pairs of sister chromatids in one nucleus (**Fig. 3d**). In CAROLI tumours, this is supported by the approximate doubling of nuclear volume (quantified from digitised histopathology, **Fig. 3e**). Some other CAROLI tumours exhibit both the mutation asymmetry of lesion segregation and approximately doubled nuclear volume (**Fig. 3e**), which could be explained by a WGD event occurring subsequently to the first division post-mutagenesis. In contrast to CAROLI, the rare symmetric tumours in the other strains do not show increased nuclear size and represent instances where both daughter lineages of an originally mutagenised cell have survived and contributed approximately equally to the sequenced tumour^17^. This illustrates profound germline differences in WGD proclivity during tumourigenesis.

Other facets of genome integrity can be interpreted from sub-chromosomal patterns of mutational asymmetry. After identifying and filtering out artefacts from reference genome misassemblies (**Extended Data Fig. 3a-d**; **Supplementary Table 4**), transitions between genomic blocks of asymmetry reveal sites of sister-chromatid exchange (SCE) that occurred in the first cell-generation after mutagenesis^14,15^. These are thought to reflect homologous repair at sites of DNA double-strand breaks, with an average 27 (s.e.m.=0.02) SCEs per tumour genome, though significantly fewer are observed in CAROLI (mean=23, s.e.m.=0.2, p=0.007, Mann-Whitney U test versus all other strains, only considering mutationally asymmetric tumours).

### Chromosome gain and loss biases are conserved

Large copy number variants (CNVs >1Mb), including whole chromosome aneuploidies, are generally rare in DEN induced tumours and most (69%) have no detectable events (**Fig. 3f**). However, CNVs are significantly enriched in CAROLI tumours with WGD (odds ratio=8.3, p=9.0×10^−7^, two-sided Fisher’s exact test, versus all other tumours), suggesting either increased tolerance of aneuploidies or greater genome instability, consistent with previous observations of WGD in cancer^26^. Chromosome 15 is recurrently duplicated in tumours from all strains and, notably, contains the *Myc* oncogene that is frequently amplified in human cancer^27^. Additionally, there is a remarkable consistency across strains for each chromosome being biassed towards either copy gain or copy loss (**Fig. 3f,g**), likely reflecting oncogenic selection for the dosage of key genes encoded within the respective chromosome. In aggregate analysis across strains, gains of chromosome 1, 2, and 15 and losses of chromosomes 4 and 19 are significantly more frequent than random expectation, representing candidate oncogenic driver events.

At multiple scales, the genetic background of each strain substantially influences genome stability following DEN treatment. This is exemplified by CAROLI, the phylogenetic out-group, which shows unique responses of frequent WGD, enrichment of CNVs, and depletion of SCEs. However, none of these mutational differences between strains obviously explain the genetic differences in tumour susceptibility. Therefore, we next identified and compared the oncogenic driver events.

### Conserved selection of driver genes

We identified genes containing candidate driver mutations using a suite of complementary driver discovery methods using 581 tumours aggregated across all strains^28–30^. Remarkably, the acquisition of activating mutations in the MAPK pathway is highly conserved across strains: 95% of tumours across all mouse strains have probable activating mutations in *Braf* (n=252), *Hras* (n=224), *Egfr* (n=84) or *Kras* (n=21) (**Fig. 4a**). Seven additional genes were under positive selection (**Fig. 4a**). Of these, *Pten*, *Crebbp*, and *Amer1* are well established tumour suppressor genes in the PI3K, Notch, and Wnt signalling pathways^31^, and *Stag2* and *Kmt2d* are tumour suppressors involved in chromatin regulation. Both *Stag2* and *Amer1* are located on the X chromosome, so their loss of function mutation was a complete knockout in the all male mice of this study. The other identified candidates are *Pyroxd2* and *Naaladl2*, neither of which are known cancer driver genes.

**Fig. 4.**
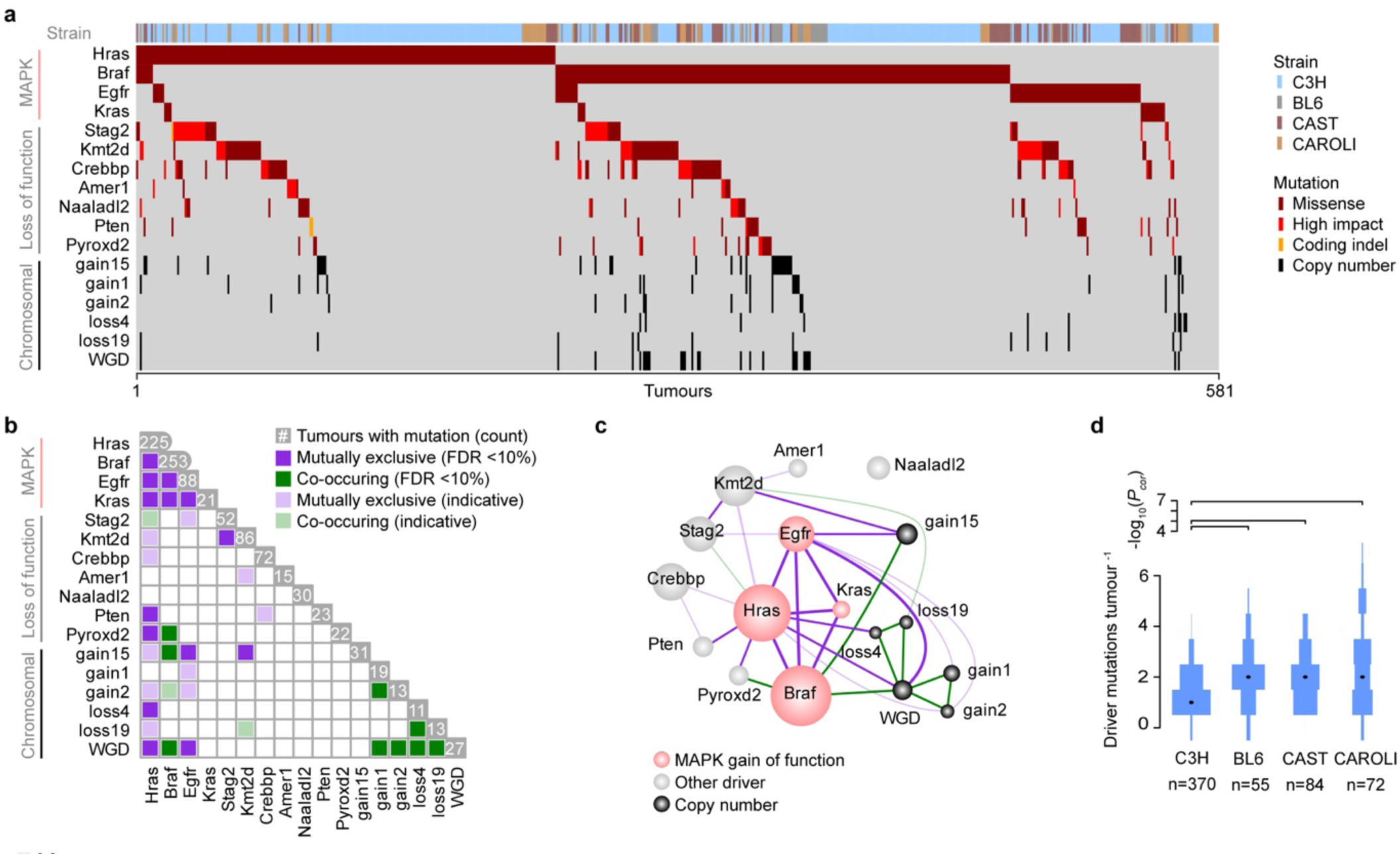
Conservation of driver genes and MAPK pathway activation. **a**, Oncoplot summarising candidate driver mutation occurrence (rows) across the 581 tumours (columns) from all strains. Reading frame disrupting substitutions such as stop-gain, stop-loss and, essential splice-site mutations are annotated as high-impact. Non-MAPK pathway driver genes frequently disrupt the reading-frame suggesting they are predominantly loss of function mutations. **b**, Significant co-occurrence (green) and mutual exclusive (purple) relationships are evident between the driver mutations, for example indicating oncogenic synergy between *Braf* activating mutations and loss of function in *Pyroxd2*. Aggregate analysis encompassing all strains. **c,** Network graph summarising the driver mutation interdependency relationships, illustrating the core oncogenic dependency on MAPK pathway gain of function mutations. The area of circles is proportional to the number of driver mutations; dependency key shared with **b**. **d**, The total number of driver mutations per-tumour systematically differs between strains (Mann-Whitney two tailed tests with Bonferroni correction).

The activating mutations in the MAPK pathway show strong mutual exclusivity that is consistently replicated across strains (**Fig. 4b,c**; **Extended Data Figs. 4 & 5**). Other candidate driver mutations show patterns of co-occurrence and mutual exclusivity that relate to the specific MAPK gene mutated, and define distinct axes^32^ of oncogenic pathway perturbation (**Fig. 4b**). These driver relationships include the significant co-occurrence of *Braf* mutations with chromosome 15 gain (gain15; weighted mutual information (wMI) p-value=0.015), the mutual exclusivity of *Stag2* and *Kmt2d* (wMI p<0.001), and a consistent trend for *Kmt2d* mutual exclusivity with *Hras* (**Fig. 4b,c**).

It is striking that in CAROLI, WGD is specific to *Braf* driven tumours (**Extended Data Fig. 4d,h**); the single WGD tumour with a *Hras* mutation also contains a *Braf* driver mutation (**Fig. 4a**). The significant co-occurrence between *Pyroxd2* and *Braf* (p=0.016) mutations, along with mutual exclusivity of *Pyroxd2* with *Hras* (wMI p=0.009), support the identification of *Pyroxd2* as a driver gene in this system (**Fig. 4b,c**).

The number of drivers per tumour also depends on background genotype. At least two driver mutations are typically identified per tumour in BL6, CAST, and CAROLI, but C3H significantly differs with a median of only one driver (p=1.97×10^−13^, Wilcoxon rank sum test test C3H versus other strains; **Fig. 4d**). This suggests that the higher liver tumour susceptibility of C3H may, in part, be attributable to fewer genetic changes being required for tumour inception than in the other strains.

In summary, despite evolutionary divergence in key aspects of mutagenesis and transformation, our novel inter-strain comparison of highly controlled tumour induction reveals remarkably ubiquitous selection of driver mutations to MAPK pathway genes. Given this convergence on a single pathway, we next examined the downstream consequences on gene regulation and the resultant tumour phenotypes.

### Consistent oncogenic perturbation of gene expression

We profiled the transcriptome of whole genome sequenced tumours (n=386) using total RNA-seq and characterised the regulatory landscape of strain-matched normal livers (P15) using ChIP-seq, ATAC-seq, and RNA-seq (**Fig. 1b**). The transcriptome analysis demonstrated strain differences in gene expression for both P15 livers and DEN induced tumours, with CAROLI, the phylogenetic outgroup, most divergent (**Fig. 5a**). We previously demonstrated that the transcription coupled repair of genes is prevalent in the DEN system^14,16,17^ and we could now examine mutation rates in the non-coding genome using experimentally-defined promoter, enhancer, and CTCF binding site maps. While there are marked differences in the mutation rate between classes of regulatory sites (bound CTCF sites > enhancers > promoters) within each strain, the ranking is common between strains (**Extended Data Fig. 6a-d**), implying that relative repair efficiencies are conserved. Next, to assess the effect of protein-DNA binding and regulatory activity on mutation rates, we identified regions that aligned across all four strains, and exhibited CTCF binding or regulatory activity in at least one strain. Aligned orthologous regions that did not show activity in a specific strain were defined as a “shadow” region in that strain (**Extended Data Fig. 6e**). Differences in mutation rate between bound and shadow regions then reveal differences in mutation properties that can be attributed to protein binding^33^. For CTCF, bound regions had higher mutation rates than unbound shadow regions which supports the finding that, in addition to impeding repair of transcription factor binding sites^34^, CTCF binding can increase DNA damage^17^. In contrast, the mutation rate of active enhancers and promoters was lower than inactive shadow regions, indicative of greater repair efficiency at the corresponding active, accessible regions^17^ (**Extended Data Fig. 6f-i**).

**Fig. 5.**
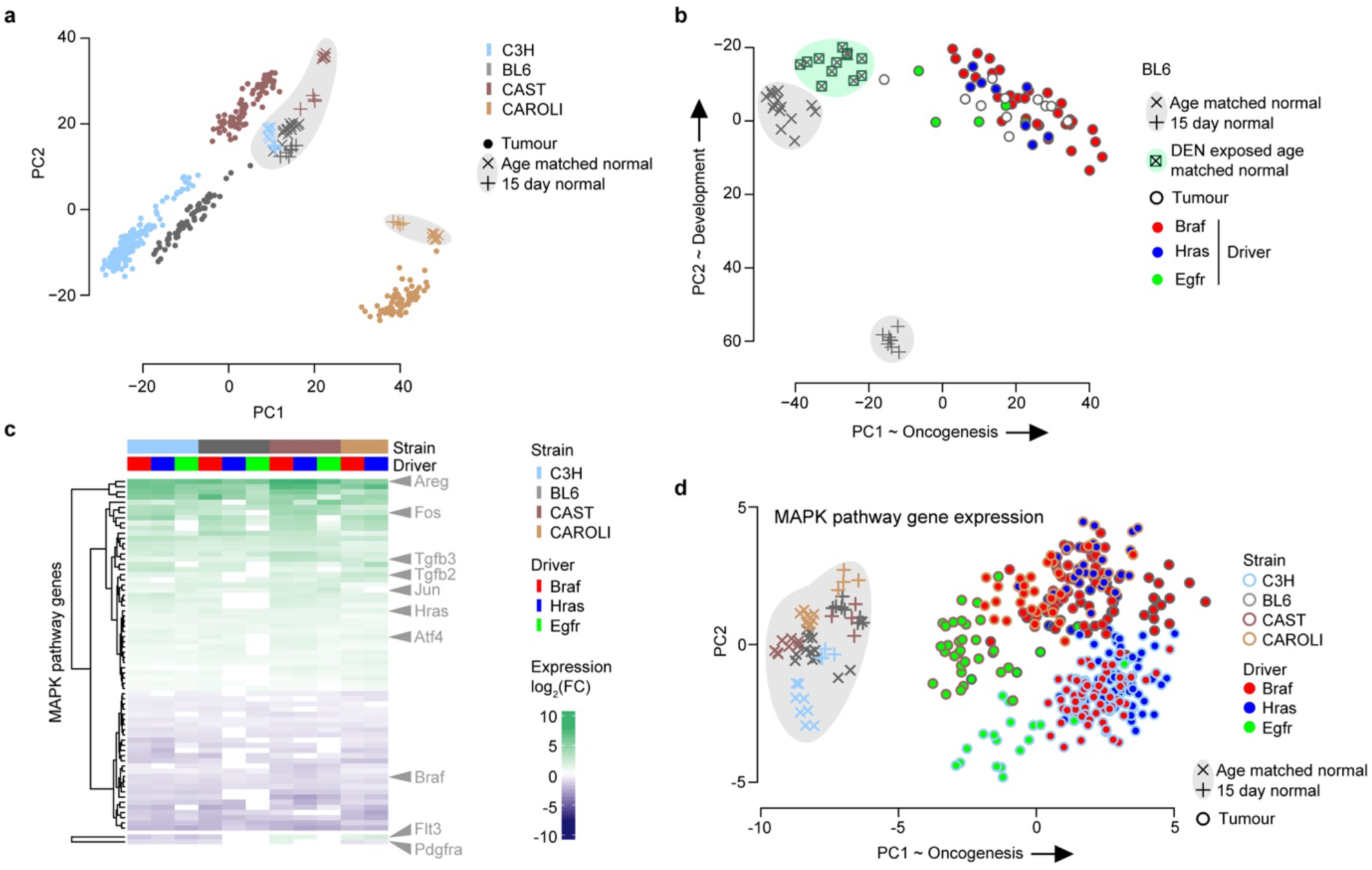
Tumours from different strains converge on a common transcriptional state. **a,** Expression analysis of DEN-induced tumours and untreated liver tissue (shaded grey; P15 and age-matched to tumour isolation) shows the first two principal components (PC1, PC2) capture expression clustering by strain (colour), developmental stage, and tumour status (shapes). **b**, Principal component expression analysis of BL6 tumours, along with comparison to background liver sampled distant to tumours in the DEN-exposed animals (green checked square), and age-matched liver tissue from DEN-naive animals (grey X). This demonstrates an axis of oncogenic transformation (PC1) that is distinct from ageing or developmental changes in gene expression (PC2). **c**, MAPK pathway genes are consistently differentially expressed between age-matched normal liver and *Braf*, *Hras* and *Egfr*-driven DEN induced tumours. The only MAPK genes with discordant, significant expression changes between strains are *Flt3* and *Pdgfra*. **d**, Principal component analysis of MAPK pathway gene expression demonstrates the axis of oncogenic transformation is consistent between strains (PC1) and highlights the muted transcriptional perturbation of *Egfr* driver mutations compared to *Braf* and *Hras*, which is also seen in **c**.

To delineate the relative contributions of development, DEN exposure, and oncogenesis, we additionally profiled the transcriptome of DEN-exposed background liver (distant to tumours) in BL6 mice (**Fig. 5b**). The majority of variance in expression between samples captures the oncogenic progression from normal liver (age-matched, untreated), through DEN-exposed background liver (non-tumour), to DEN-induced tumour (PC1, 50% of variance). In contrast, the second principal component corresponds to developmental time from P15 to adult gene expression (18% of variance). For genes that are differentially regulated between age-matched normal liver and DEN induced tumours, there is a high degree of concordance in both the direction and magnitude of expression changes when compared between strains (95-99% pairwise concordance; **Extended Data Fig. 7**). Intriguingly, for long non-coding RNA transcripts that are differentially expressed, there is a pronounced bias towards their reduced expression in tumour compared to age-matched normal (**Extended Data Fig. 7**).

Driver mutation analysis highlighted the MAPK pathway as central to oncogenic transformation in each strain. We find that many genes in the MAPK pathway are differentially expressed between age-matched normal and DEN induced tumour, with the tumour associated changes in expression consistent between strains (**Fig. 5c**). Principal component analysis of MAPK pathway gene expression separates tumour from non-tumour samples on PC1 for all strains, though *Egfr* mutation driven tumours cluster closer to normal liver than *Braf* or *Hras* mutation driven tumours (**Fig. 5d**). Remarkably, the differences between the phylogenetic outgroup, CAROLI, and the other strains are absent in the expression patterns observed in MAPK genes. In summary, different driver mutations in MAPK genes arising in the context of different genetic backgrounds all converge on a strikingly consistent, tumour-associated transcriptional phenotype.

### Genetic background strongly biases driver utilisation

Despite almost universal acquisition of activating mutations in the MAPK pathway (**Fig. 4a**) and cross-strain convergence of MAPK expression perturbation (**Fig. 5c**), the frequency with which each MAPK driver gene is mutated varies significantly between strains (χ^2^ p=6.59×10^−14^; **Fig. 6a**). Even more remarkably, although mutations at codon 61 of *Hras* are found as prominent oncogenic drivers in all four strains (and among human cancers^35^), the identity of the specific oncogenic amino acid change is highly divergent between genetic backgrounds (χ^2^ p=2.69×10^−6^; **Fig. 6a**).

**Fig. 6.**
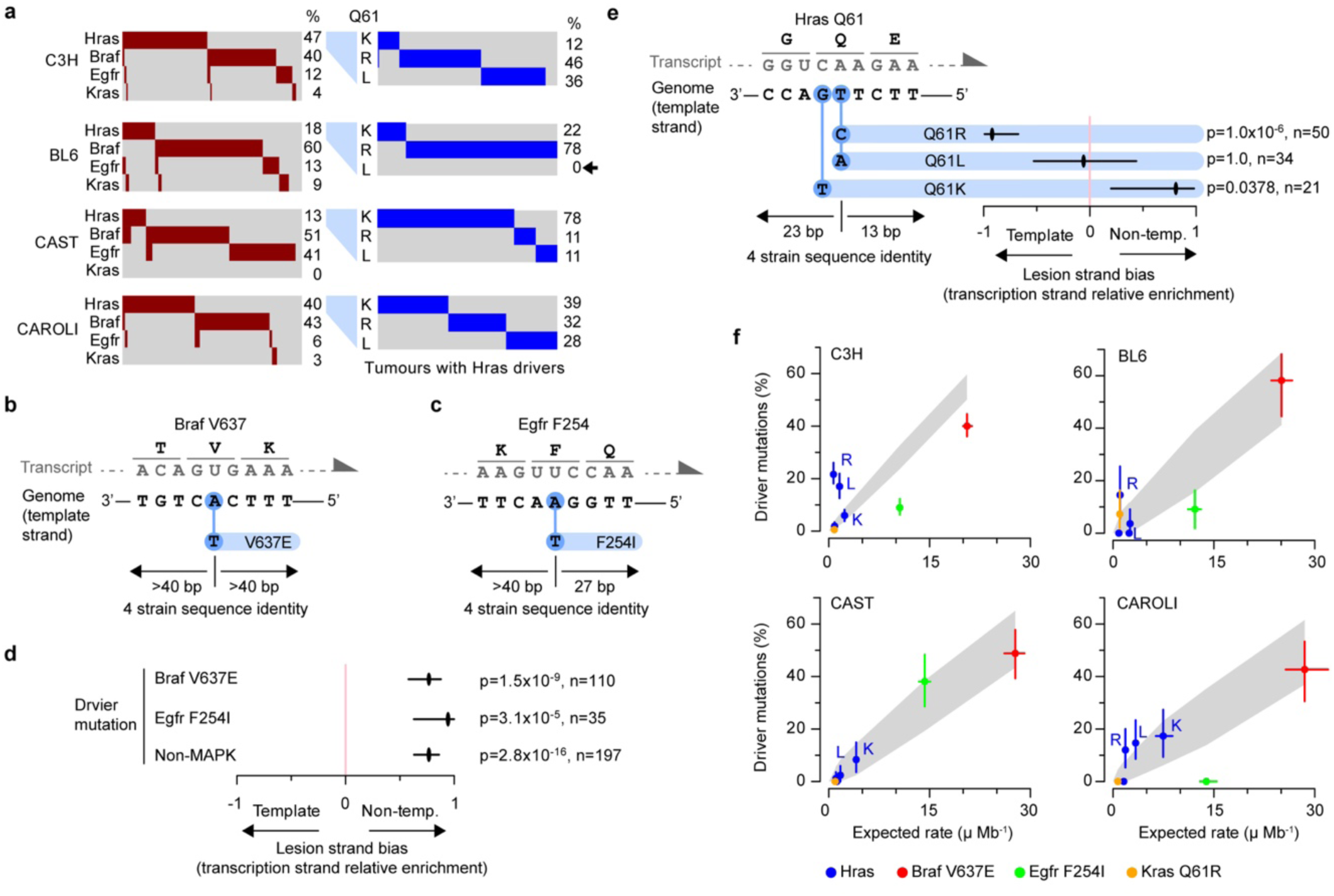
Genetic background influences the identity of MAPK pathway activating mutations. **a,** Most tumours have a MAPK pathway activating mutation, but there are strain differences in the proportion of the mutations in each gene (left panels). In tumours with *Hras* driver mutations (right panels), there are profound strain differences in the specific mutation found at the Q61 codon, notably including the complete absence of Q61L in BL6 (arrow). **b,** Schematic of the *Braf* V637E driver mutation: An adenosine (A) nucleotide on the genic template strand is mutated to thymine (T). The immediate sequence context is identical between all four strains for at least 40 nucleotides both upstream and downstream of the mutated site. **c,** Summary of the *Egfr* F254I driver mutation (as **b**)**. d,** There is a strong preference for recurrent *Braf* and *Egfr* driver mutations to be found in tumours that inherited DNA damage (lesions) on the non-template strand (not subject to transcription coupled repair) based on lesion strand bias (Methods; aggregate analysis over all 4 strains, whiskers show 95% CI from bootstrap sampling). Aggregate analysis over all non-MAPK pathway driver mutations shows the same bias for non-template strand damage. Lesion strand bias statistical tests are two-sided Fisher’s exact tests; p-values shown after Bonferroni correction for multiple testing. **e,** Summary of *Hras* driver mutations (as **b,c**). *Hras* Q61K mutations (codon position 1) show non-template strand bias, concordant with other drivers, whereas mutations at the *Hras* Q61 codon position 2 (Q61L and Q61R) do not show this non-template strand damage bias. **f,** Observed percentage of tumours with each recurrent MAPK activating mutation (y-axes) compared to the expected mutation rate (μ, x-axes). Expected mutation rates were calculated based on lesion-strand resolved mutation rates and spectra, adjusting for transcription template strand and gene expression level (Methods). Confidence intervals (95%, whiskers) for both observed driver mutation percentage and expected mutation rate were calculated from bootstrap sampling of tumours. Annotations (R, L, K; blue) identify specific amino acid substitutions of *Hras* Q61. Grey areas show the predicted position (95% confidence interval) of driver mutations if their relative observed frequency depended only on the expected mutation rates.

The extended sequence context of each MAPK driver mutation is highly conserved, with no germline genetic differences within at least 13 nucleotides in any strain (**Fig. 6b-e**), thus excluding proximal sequence divergence as the explanation for different driver mutations. With the exception of *Hras* Q61R and Q61L mutations, we find that all MAPK and other identified driver base substitutions are strongly biased to have been produced from DEN induced damage on the non-template, rather than transcription template, strand (**Fig. 6d,e**; **Extended Data Fig. 8a**). This likely reflects the high efficiency of transcription coupled repair of base alkylation damage on the template strand^16,17^. An important implication of this is that, since the template strand is unaffected, mutated oncogenic proteins would not be produced until after the first round of replication following damage. In contrast the *Hras* Q61R mutations arise from template strand damage and could lead to the production of pro-oncogenic miscoded proteins before replication^36^.

To further investigate the causes of strain divergence in MAPK driver mutation utilisation we modelled the expected rate of driver mutation occurrence (**Fig. 6f**). This modelling accounted for heterogeneity of mutation burden and mutation spectrum between tumours, and captured the pronounced effects of transcription coupled repair on mutation rate (**Extended Data Fig. 8b,c**). The rank-order of expected MAPK mutations is closely matched between strains, and subtle differences in mutation spectra do not explain the dramatic phylogenetic divergence in driver utilisation (**Fig. 6f**).

Both the *Hras* Q61R and Q61L mutations are observed at higher frequencies than expected from modelling in several strains (**Fig. 6f**). Mutation spectrum suggests, and lesion strand bias confirms, that Q61R arises from T damage on the transcription template strand (**Fig. 6e**; **Extended Data Fig. 8a**). The low expected rates for *Hras* Q61R and Q61L mutations reflect the high efficiency of transcription coupled repair of DEN damaged nucleotides^14,16^. Modelling shows that the observed driver mutation frequencies could be better explained if *Hras* is not transcribed in tumour progenitor cells during the time between DNA damage and the first round of DNA replication (**Extended Data Fig. 8b,c**). This would be consistent with transformation occurring in a minority population of hepatocytes that are transiently quiescent at the *Hras* locus, which would be masked by bulk transcriptomic analysis.

Despite *Hras* Q61L and Q61R representing alternate changes of the same nucleotide that occur with similar frequency to each other in all other strains, Q61L is entirely absent from BL6 tumours (**Fig. 6a**). This suggests genetic background differences in oncogenic selection. A potential source of such differences is neoepitope presentation in the major histocompatibility (MHC) class I molecules. The *Hras* Q61L mutation is predicted to have high affinity for MHC class I uniquely in BL6 (**Extended Data Fig. 8d**), possibly explaining the strain specific absence of this mutation through epitope presentation and immune clearance. However, predictions of neoepitope presentation do not explain all strain differences in driver mutation frequency, and immune editing does not appear to be a major force shaping the observed distribution of mutations (**Extended Data Fig. 8e,f**). We next considered more general strain differences of within-tumour oncogenic selection.

### Genetic background shapes oncogenic selection

The persistence of DNA damage through multiple rounds of replication can generate multiallelic variation - multiple different mutations at the same site - within an expanding clone^14,15,37^. This multiallelic variation represents the generation of extensive genetic diversity within the first few cell divisions following a burst of mutagenesis, affecting both the presence or absence of subclonal mutations, and their co-occurrence within a sublineage. The segmental pattern of multiallelic variation across the tumour genome can identify which cell generation post-mutagenesis gave rise to that tumour’s most recent common ancestor (MRCA)^14^. If a tumour’s MRCA is a first-generation daughter of a mutagenised cell, then one strand of every chromosome retains DNA damage and results in genome-wide multiallelic variation. Damage-containing strands randomly segregate through subsequent cellular generations, losing on average 50% of damaged strands per generation. Consequently, a tumour developing from a second generation MRCA is expected to have only 50% of the genome as multiallelic, 25% in a third generation MRCA, and 12.5% in a fourth generation MRCA (**Fig. 7a**).

**Fig. 7.**
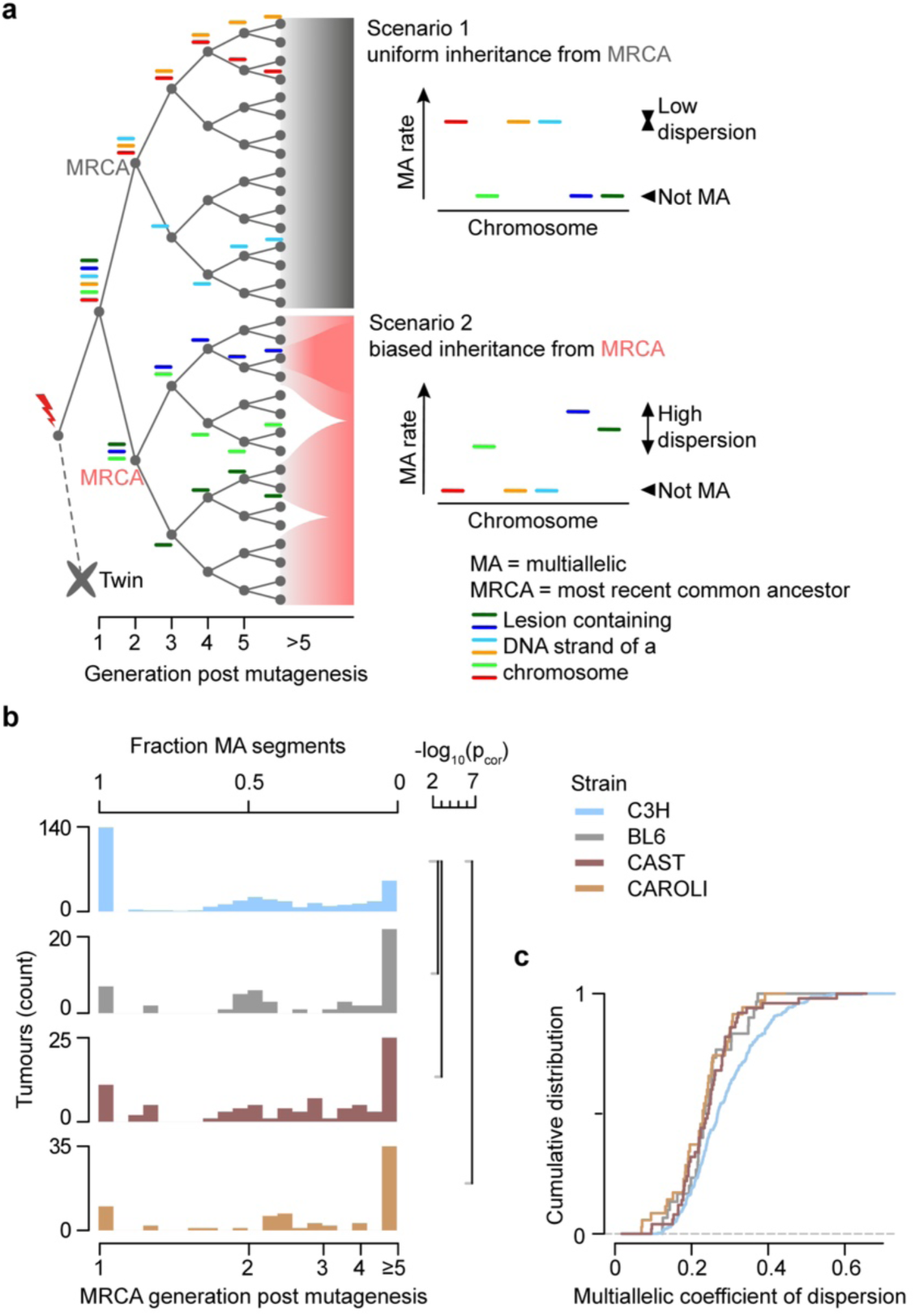
Genetic background variation impacts tumour clonal dynamics. **a,** Schematic illustrating the segregation of lesion-containing chromosome strands (coloured bars) through cell generations (x-axis) following mutagenesis (red bolt). Typically only one daughter of an originally mutagenised cell contributes to a tumour, so the twin lineage is lost (dashed line). Scenario 1 (upper panel, grey) shows a tumour that develops from a most recent common ancestor (MRCA) in the second generation, with all subsequent lineages contributing equally. Chromosomes without a lesion strand in the MRCA are not multiallelic (MA) in the tumour; chromosomes with a lesion strand in the MRCA all have approximately the same MA rate in the tumour. Scenario 2 (lower panel, pink) also shows the evolution of a tumour with a second generation MRCA, but subsequent lineages have unevenly contributed to the tumour resulting in heterogeneity of MA rate between chromosomes that inherit a lesion containing strand. **b,** The fraction of chromosomal segments with multiallelic variation (x-axis) identifies the cell-generation post mutagenesis of each tumour’s MRCA. The frequency distribution (y-axis, tumour count) of MRCA generation is strikingly different between C3H and the other strains (two-sided Kolmogorov-Smirnov test p-values, Bonferroni corrected, shown for pairwise comparisons p<0.05), with the majority of C3H tumours having their MRCA in the first cell generation post-mutagenesis. **c,** Compared to the other strains, C3H (blue, key shared with panel **b**) has a greater proportion of tumours (cumulative distribution, y-axis) with a high multiallelic coefficient of dispersion (x-axis), indicating unequal contributions of sub-clones to the sequenced tumour cell population.

Quantifying multiallelic rates across the tumour genomes demonstrates that all strains are able to instantly transform; that is, where a first generation post-mutagenesis cell is the MRCA of a tumour (**Fig. 7b,c**, **Extended Data Fig. 9a**). This is remarkable because the first generation MRCA tumours have essentially retained all the multiallelic and combinatorial genetic diversity generated from the initial burst of DNA damage and exhibit minimal loss of sublineages. Instant transformation is consistent with aspects of the big bang model of transformation^38^, but contrasts with a prevalent view of cancers developing through strong selection and recurrent subclonal sweeps^39,40^. Notably, C3H represents a clear outlier, with greater propensity for a first generation MRCA than any of the other strains, implying reduced selective pressure on early tumours in C3H (two-sided Kolmogorov-Smirnov test, C3H versus the aggregate of other strains, p=6.6×10^−9^; **Fig. 7c**).

By considering the heterogeneity of multiallelic rates across a tumour genome we can estimate ongoing (more recently than the MRCA) subclonal selection within a tumour (**Fig. 7a**). Tumours from all four strains exhibit similar distributions of post-MRCA selection (**Extended Data Fig. 9b-e**), but a subset of C3H tumours (19% of analysable, using non-C3H as null distribution) show evidence of higher subclonal, post-MRCA selection, reiterating that germline background (strain) influences oncogenic selection.

## Discussion

Despite decades of human cancer genome sequencing, how an individual’s genetic background shapes the initiation and early progression of tumours remains poorly understood. A major constraint has been our inability to rigorously control for variables inherent in human life, including diet, sex, environment, mutagen exposure, demographic history. We reasoned that a well established mouse model of liver cancer would control these variables, and thus enable the systematic interrogation of genetic background effects on early cancer development^9,10,13,14^. By characterising cancer development in four inbred mouse strains, whose phylogenetic divergence mimics and extends beyond that of the human population^23–25^, we revealed the extent to which genetic background influences both mutational processes and the paths to tumour development.

On one hand, our comparative analyses of these tumour cohorts show features that are largely independent of genetic background. The mutational loads and spectra created by identical mutagenic treatments are highly similar between lineages. Aneuploidies are generally rare, but when they do occur specific chromosomes show pronounced biases towards amplification or deletion that are conserved between lineages. In addition, the acquisition of DEN induced activating mutations in the MAPK pathway genes *Braf*, *Hras*, *Egfr* or *Kras* was an almost universal feature, reflecting their prevalence in human pan-cancer analyses^41,42^. Regardless of genetic background or the specific driver mutations, the tumours in our study all acquired similar transcriptional perturbation of the wider MAPK pathway: This perturbation is also seen in up to 50% of human hepatocellular carcinomas^43^ and almost all with advanced stage disease^44^, suggesting that tumour development converges on a common regulatory state.

On the other hand, this highly controlled system revealed three striking phenotypes dependent on genetic background. First, despite the commonality of MAPK activating mutations, there were pronounced differences in the occurrence of specific activating mutations between strains. For example, *Hras* Q61L is common in the tumours of C3H (17%) but not found in BL6 tumours, whereas *Hras* Q61R, a different change to the same nucleotide is common in both strains. These differences cannot be explained by subtle divergence in mutation spectra between lineages, but instead point to differences in selection imposed by the germline genome. In the case of *Hras* Q61 mutations, differences in epitope presentation and immune editing may account for the absence of specific alterations in BL6, but alternate mechanistic explanations are required for other biases in driver usage, such as *Egfr* F254I between CAST and CAROLI.

Second, genetic background profoundly influences the intensity of oncogenic selection. For instance, tumours in most strains show the extensive loss of sub-lineages, with the sequenced tumours containing only a fraction of the genetic diversity generated from an initial burst of mutagenesis. In contrast, on the C3H genetic background there are minimal lineage losses and most of the initially generated genetic diversity is typically retained in the macroscopic tumour reflecting the retention of all sub-lineages of an originally mutagenised cell. Furthermore, genetic background differences in the number of driver mutations identified per tumour suggest that C3H requires fewer genetic changes to achieve oncogenic transformation, or tumour survival, than the other strains. These observations of minimal clonal selection in C3H align with the short latency, rapid growth, high tumour burden, and high spontaneous cancer susceptibility of this genetic background, establishing a link between within-tumour selection and germline cancer susceptibility.

Third, we found that genetic background has a major influence on the occurrence of mononuclear whole genome duplication (WGD), which was seen exclusively in CAROLI tumours. From their mutational symmetry, these WGD events must have occurred in the first cell cycle following mutagenesis and always co-occur with *Braf* driver mutations. WGD might be a pro-oncogenic event in these tumours that synergises with, or is caused by, *Braf* activating mutations. Alternatively, the WGD could occur more widely in response to DNA damage in CAROLI, but only those with a *Braf* driver mutation being able to develop into tumours. This latter scenario could be consistent with the titration of MAPK pathway signalling to promote cell growth while avoiding oncogene induced senescence^45^. Whether WGD in response to DNA damage is initially pro-oncogenic or not, once it has occurred the risk of secondary aneuploidies is dramatically increased, providing new opportunities for oncogenic selection. For many patients WGD is a critical, often early^46^, event in tumour evolution that confers poor prognosis^47–49^. Recent observations of elevated WGD rates in cancers from specific ethnic groups^50^ support our conclusion that germline variation influences the risk of this pivotal change during cancer development.

Importantly, our finding that genetic background impacts both mutation and selection during tumourigenesis, despite identical exposures, tells us that within the human population, an individual’s genetic background is likely to interact with both (i) how a given environmental exposure initiates and promotes tumourigenesis and (ii) the efficacy and toxicity of subsequent genotoxic cancer treatments. This may help to explain differences in molecular profiles of cancers arising in populations of different ancestries^51^ and clinical outcomes between patient groups^50,52^. In this context, it also raises challenging questions about how we should take genetic background into account in identifying higher risk groups for screening, pre-invasive disease surveillance, and identifying prognostic and predictive biomarkers.

Building on decades of epidemiological studies^50,51,53–59^, we now provide compelling *in vivo* evidence that subtle germline variation significantly and reproducibly impacts multiple aspects of tumourigenesis. These are important precedents to establish because the success (or failure) of many research efforts and clinical trials is ultimately determined by how well the evolutionary trajectory of cancers can be pre-empted. Our findings highlight the importance of considering genetic background as an important variable in the design and interpretation of biomedical and translational research.

## Methods

### Mouse colony management

Animal experimentation was carried out in accordance with the Animals (Scientific Procedures) Act 1986 (United Kingdom) and with the approval of the Cancer Research UK Cambridge Institute Animal Welfare and Ethical Review Body (AWERB): the maximum approved tumour burden was 10% body weight, which was not exceeded. Animals were maintained using standard husbandry: mice were group-housed in Tecniplast GM500 IVC cages with a 12 h:12 h light:dark cycle and *ad libitum* access to water, food (LabDiet 5058), and environmental enrichments.

The following mouse strains and species were used: *Mus musculus domesticus* C3H/HeOuJ (hereafter referred to as C3H mice) and C57BL/6J (BL6), *Mus musculus castaneus* CAST/EiJ (CAST), and *Mus Caroli* CAROLI/EiJ (CAROLI) (**Supplementary Table 2**). For simplicity, these strains, subspecies, and species are herein referred to as ‘strains’.

### Chemical model of hepatocarcinogenesis

15-day-old (P15) male mice of all strains were treated with a single intraperitoneal (IP) injection of N-Nitrosodiethylamine (DEN; Sigma-Aldrich N0258; 20 mg/kg body weight) diluted in 0.85% saline. Liver tumour samples were collected from DEN-treated mice 25 weeks (C3H), 36 weeks (BL6), 38 weeks (CAST), or 78 weeks (CAROLI) after treatment; pilot data indicated that 100% mice would develop tumours by these timepoints. All macroscopically identified tumours were isolated and processed in parallel for DNA and RNA extraction and histopathological examination; background non-tumour liver was also sampled from BL6 mice. Untreated mice from each strain were assessed for the presence or absence of tumours at these timepoints, and also aged further to assess the inherent susceptibility to spontaneous tumours (**Supplementary Table 3**). Additional tissues from untreated P15 mice (ear, tail, and liver) and untreated age-matched adult mice (liver only; C3H 27 weeks, BL6 38 weeks, CAST 40 weeks, CAROLI 80 weeks) were sampled for control experiments.

### Tissue collection and processing

Liver tumours of sufficient size (≥2 mm diameter) were bisected; one half was flash frozen in liquid nitrogen and stored at −80°C for DNA and RNA isolation, and the other half was processed for histology. Tissue samples for histology were fixed in 10% neutral buffered formalin for 24 h, transferred to 70% ethanol, machine processed (Leica ASP300 Tissue Processor; Leica, Wetzlar, Germany), and paraffin embedded. All formalin-fixed paraffin-embedded (FFPE) sections were 3 μm in thickness.

### Histochemical staining

FFPE tissue sections were haematoxylin and eosin (H&E) stained using standard laboratory techniques. Histochemical staining was performed using the automated Leica ST5020; mounting was performed on the Leica CV5030.

### Whole slide image acquisition

Tissue sections were digitised using the Aperio XT system (Leica Biosystems) at 20x resolution; all H&E images are available in the BioStudies archive at EMBL-EBI under accession S-BSST383^14^ and S-BSST384 (this study).

### Tumour histopathology

H&E sections of liver tumours were blinded and assessed twice by a histopathologist (S.J.A). Tumours were classified according to the International Harmonization of Nomenclature and Diagnostic Criteria for Lesions in Rats and Mice (INHAND) guidelines^61^. In addition, tumour grade, size, morphological subtype, nature of steatosis, and mitotic index were assessed (**Supplementary Table 1**), as well as the presence of cystic change, haemorrhage, necrosis, or vascular invasion.

### Sample selection for computational pathology, WGS, and RNA-seq

Tumours which met the following histological criteria were selected for whole genome sequencing (C3H n=370, BL6 n=55, CAST n=84, CAROLI n=72): (i) diagnosis of dysplastic nodule (DN), (ii) homogenous tumour morphology, (iii) tumour cell percentage >70%, and (iv) adequate tissue for DNA/RNA extraction. Neoplasms with extensive necrosis, mixed tumour types, a nodule-in-nodule appearance (indicative of a hepatocellular carcinoma (HCC) arising within a DN), or contamination by normal liver tissue were excluded. Since carcinogen-induced tumours arising in the same liver are independent^13^, multiple tumours were selected from each mouse to minimise the number of animals used. A subset of normal (non-tumour) samples from untreated age-matched mice were also whole-genome sequenced (C3H n=13, BL6 n=11, CAST n=7, CAROLI n=14). In addition, a small number of HCC tumours were subject to WGS and released in the accompanying data, but were excluded from the analyses reported in this manuscript (C3H n=1, BL6 n=11, CAST n=0, CAROLI n=4).

### Computational pathology analysis

Whole slide images (WSIs) of tumours were annotated in QuPath (v.0.2.2)^62^ using the polygon tool to include neoplastic tissue and exclude adjacent parenchyma, cyst cavities, processing artefacts, and white space. For tumours with multiple transections, only a single WSI was used. Annotations were reviewed for quality by two histopathologists (S.J.A. and J.C.). Annotated regions were tessellated into fixed size, non-overlapping 256 µm x 256 µm tiles using Groovy in QuPath. For segmentation of epithelioid nuclei, a pre-trained StarDist^63^ model (he_heavy_augment.zip) was downloaded from https://github.com/stardist/stardist-imagej/tree/master/src/main/resources/models/2D and an inference instance was deployed using Groovy across the tiles in QuPath, built from source with Tensorflow^64^, with a minimum detection threshold of 0.5. Quantitative geometric features (size and shape) were measured for each nuclear object (**Supplementary Table 1**). The object-level measurements were subsequently abstracted by computing the median, standard deviation, interquartile range, and kurtosis for each tile and WSI. Nuclear volume was subsequently calculated using the median nuclear diameter per slide.

Digitised histology images of DEN-induced tumours are available from Biostudies: Accession numbers S-BSST383 https://www.ebi.ac.uk/biostudies/studies/S-BSST383 and S-BSST384 https://www.ebi.ac.uk/biostudies/bioimages/studies/S-BSST384.

### DNA and RNA isolation

Simultaneous isolation and purification of genomic DNA and total RNA from tumours and background liver tissue was performed using the AllPrep 96 DNA/RNA Kits (Qiagen 80311), according to the manufacturer’s instructions. DNA from P15 liver tissues was isolated and purified using the AllPrep DNA/RNA mini kit (Qiagen 80204); total RNA was extracted using QIAzol Lysis Reagent (Qiagen 79306), according to the manufacturer’s instructions. Genomic DNA was isolated from ear/tail samples using the DNeasy blood & tissue kit (Qiagen 69504), according to the manufacturer’s instructions.

### Whole genome sequencing (WGS)

Genomic DNA quality was assessed on a 1% agarose gel and quantified using the Quant-IT dsDNA Broad Range Kit (Thermo Fisher Scientific Q33130). Genomic DNA was sheared using a Covaris LE220 focused-ultrasonicator to a 450 bp mean insert size.

WGS libraries were generated from 1 μg of 50 ng/ul high molecular weight gDNA with a *S. cerevisiae* spike-in control (1 ng of 900 pg/ul genomic DNA; Merck Millipore 69240) using the TruSeq PCR-free Library Prep Kit (Illumina 20015963), according to the manufacturer’s instructions. Library fragment size was determined using a Caliper GX Touch with a HT DNA 1k/12K/Hi Sensitivity LabChip and HT DNA Hi Sensitivity Reagent Kit to ensure 300-800 bp (target ∼450 bp).

Libraries were quantified by real-time PCR using the Kapa library quantification kit (Kapa Biosystems KK4824) on a Roche LightCycler 480. 0.75 nM libraries were pooled in 6-plex and sequenced on a HiSeq X Ten (Illumina) to produce paired-end 150 bp reads. Each pool of 6 libraries was sequenced over eight lanes (minimum of 40x coverage).

WGS FASTQ files are available in the European Nucleotide Archive at EMBL-EBI under accession PRJEB37808^14^ and PRJEB15138 (this study).

### Whole genome sequencing read alignment

Alignment of WGS data were performed as described^14^. In brief, sequencing reads were aligned to their respective genome assemblies from Ensembl v.91^65^, (BL6 = GRCm38 = GenBank: GCF_000001635.26, C3H = C3H_HeJ_v1 = GenBank: GCA_001632575.1, CAST = CAST_EiJ_v1 = GenBank: GCA_001624445.1, CAROLI = CAROLI_EiJ_v1.1 = GenBank: GCF_900094665.2) using bwa-mem (v0.7.12)^66^. The yeast reference genome S228c (GenBank: GCA_902192305.1) was included in the alignment targets to account for the spike-in DNA. Using WGS of non-tumour liver, ear and tail samples (described above) collected and sequenced contemporaneously with tumour samples, regions of abnormal WGS read coverage were identified^14^ and subsequently masked from the analysis (percentage of the reference genome masked: BL6 = 5.5%, C3H = 12.7%, CAST = 11.5%, CAROLI = 12.5%).

### Variant calling and mutation filtering

Single base substitution (SBS) mutations were called using Strelka2 (v.2.8.4)^67^. As described in^14^, fixed genetic differences and segregating germline variants within our colonies were identified and discarded. Insertion and deletion (collectively indel) mutations were filtered with identical parameters to SBS mutations with the additional criteria that (i) each called indel mutation must be supported by at least 3 independent reads, and (ii) where the same indel mutation was found in at least two tumours from the same animal and shared between >50% of tumours from that animal it was filtered from the calls of all tumours within the strain. This latter “animal-level-filtering” step removes fixed and segregating germline indels that passed the familial clustering filter due to the conservative calling of indels relative to SBS mutations. Of the 4,446 indel mutations subject to animal-level-filtering (distinct sites, not summing repeat occurrences over tumours), only 2 are predicted by Variant Effect Predictor (VEP; v.109.3)^68^ to disrupt protein sequence (genes *Atxn2l* and *Cyp4a30b*).

### Mutation rate calculations

For lesion-strand resolved analyses, SBS mutation rates were calculated as 192 category vectors representing every possible single nucleotide substitution conditioned on the identity of the upstream and downstream nucleotides. Each rate being the observed count of a mutation category divided by the count of the trinucleotide context in the analysed sequence. For lesion-strand non-resolved analyses, the same procedure was followed but reverse complement mutation (e.g. T→C and A→G) and sequence context (e.g. ATG and CAT) counts were combined to give 96 category vectors, which by convention are presented oriented as a mutation from the pyrimidine base of the basepair. Indel rates were calculated as indel count per megabase, per tumour and corrected for the expected diploidy of autosomes.

### Mutation signature analysis

Single base substitution (SBS) mutation signatures were defined and deconvolved using the R NMF library (v.0.28) nmf function^69^ with rank=5, run=1,500 and the method set to use the Brunet algorithm^70^. The rank defines the number of signatures to identify, the value of five was selected after testing ranks 2 to 8 inclusive and selecting the first value of rank for which the cophenetic coefficient substantially decreases^70^. This value also coincided with an inflection point in the curve plotting the residual sum of squares against rank, as proposed as an alternate method for optimal rank choice^71^.

Indel mutation signatures were generated using SigProfiler (1.2.19)^72^ using the corresponding strain specific reference assemblies (masked, see above). Bootstrap replicate datasets were generated using the SigProfilerMatrixGenerator for the calculation of 95% confidence intervals. For the analysis of multi-nucleotide indel sequence composition, only tumours with cellularity estimated at >50% were considered and indels of length greater than 1 bp were folded into unique discrete sequence categories combining repeat unit repetition and reverse-complement relationships, e.g. AC deletion encompasses AC, CA, GT and TG deletion. Insertions and deletions were categorised by length (span of deleted or inserted nucleotides) and the mutation rates for each sequence category (e.g. AC) calculated as the number of events (e.g. AC deletions) divided by the number of that sequence context in the reference genome (e.g. AC occurrences). Percentage rates were calculated (e.g. AC deletion rate as a percent of the sum of dinucleotide category deletion rates) and compared between strains (**Extended Data Fig. 2e-g**); the comparison of rates thus controls for the minor sequence composition differences between the genomes of the four strains. The 95% confidence intervals were obtained from 100 bootstrap samples of the tumours for analysis within each strain.

### Mutational asymmetry segmentation and scoring

The lesion segregation model predicts whole chromosomes would be coherently strand asymmetric for mutations following a burst of DNA damage. While this is often the case, homologous repair in the first round of DNA replication following damage can result in sister-chromatid exchange events and discrete transitions in the mutational asymmetry of the chromosome such that chromosomes are made up of multi-megabase blocks of alternating mutation asymmetry^14,15^. We refer to these blocks as genomic segments. Genomic segmentation on mutational asymmetry was performed as previously reported^14^. Mutational strand asymmetry was scored for each genomic segment using the relative difference metric *S=*(*R_T_-R_A_*)*/*(*R_T_+R_A_*) where *R_T_* is the rate of mutations from T on the forward (plus) strand of the reference genome and *R_A_* is the rate of mutations from A on the plus strand, equivalent to the rate of mutations from T on the reverse (minus) strand.

### Identifying and filtering reference genome mis-assemblies

Since lesion segregation, mutation asymmetry patterns allow the long-range phasing of chromosome strands, they can detect discrepancies in sequence order and orientation between the sequenced genomes and the reference. We identified autosomal asymmetry segments that immediately transitioned from forward strand bias (*S* > 0.33) to reverse strand (*S* < −0.33) or vice versa without occupying the intermediate unbiased state (−0.33 ≥ *S* ≤ 0.33); such discordant segments are unexpected. Allowing for ±100 kb uncertainty in the position of each exchange site we produced the discordant segment coverage metric. At sites with discordant segment coverage >1 we calculated relative enrichment for mis-assembly *M=(ds-cs)/(ds+cs)* where *ds* is the number of discordant segments over the exchange site and *cs* the number of concordant: where either forward or reverse mutational asymmetry extends at least 1×10^6^ nucleotides on both sides of the exchange site. Values of *M* > 0 indicate consensus for misassembly. The approximate genomic coordinates for a C3H strain specific inversion on chromosome 6 were previously reported^73^.

### Classification of mutational asymmetry

Based upon previously described mutational asymmetry segmentation and scoring, samples were classified based on their adherence to the expectations of lesion segregation, eg: asymmetric or non-asymmetric. Firstly to reduce noise and minimise the influence of large CNVs and aneuploidies, genomic segments with <100 mutations and those with significantly higher or lower than average read depth were removed (p<0.05, probability of each segment being sampled from a normal distribution with mean and standard deviation equal to the weighted mean and standard deviation of the per-segment means). Genomic segments were classified as symmetric (abs(*S*) < 0.2, where ‘abs(*S*)’ is the absolute value of the previously defined mutational asymmetry parameter: ‘*S*’), asymmetric (abs(*S*) > 0.8) or intermediate (abs(S) ≥ 0.2 and ≤ 0.8). For each tumour, the proportion of the genome belonging to each class was calculated. Samples with <10% asymmetric autosomes were defined as non-asymmetric (symmetric). All remaining samples were classified as asymmetric. As sister chromatid exchange events can only be identified in the presence of mutation asymmetry, the non-asymmetric tumours were excluded from sister chromatid exchange rate calculations.

### Tumour cellularity estimates

As described in^14^, variant allele frequency (VAF) was calculated as (1 − *R*/*d*), where *R* is the reference read count at a mutated site and *d* is the total read depth at the site. For each tumour, the modal VAF was determined using the function ‘amps’ from the R package modes (0.7.0) to detect the size and position of the largest peak in the VAF density distribution. Cellularity was then calculated as ploidy × the major peak VAF, in which ploidy was set as 2 for all samples, except CAROLI tumours with no mutational asymmetry which were inferred to be whole-genome duplicated (ploidy = 4).

### Large copy number variant calling

Tumour genomes were segmented based on their change in read depth, relative to untreated adult liver samples (BL6, n=11; C3H, n=11; CAROLI, n=14 and CAST, n=7), using CNVkit (v0.9.6)^74^. To suppress calling of small CNVs, minimum segment size was set as 1Mb. For each segment, copy number (CN) was calculated using the log_2_ read depth fold change (FC) given by CNVkit, and the sample ploidy (p) and cellularity (c), where: CN = ((2^FC^-1+c) / c) × p. Within strain, CN estimates were adjusted for read depth biases by subtracting the median change in CN of all overlapping segments (overlapping defined as segments where the intersect is ≥80% of the union, adjustment performed for segments with ≥3 overlapping segments, change in CN calculated as CN-p). Segments with adjusted CN (adj.CN) within 0.15 of the sample ploidy were discarded. Segments with adj.CN within 0.15 an integer were called as clonal CNVs. Segments outside this threshold were identified as subclonal CNVs. Filtering was then performed to remove recurrent artefactual CNVs by discarding any short CNV (<10% of total chromosomal length) with ≥80% overlap of another CNV from the same strain.

Aneuploidies were identified as either gain or loss of chromosomal segments totalling >80% of a chromosome; typically these were whole chromosome (100%) gains and losses. Significant recurrence was identified by randomly permuting aneuploidy labels across autosomes. P-values were calculated by comparing the observed recurrence of a specific aneuploidy to the null distribution of random recurrence generated by 10,000 permutations. Aneuploidies with p<0.05 Bonferroni corrected significance were identified as having significant recurrence.

### Multiallelic mutation rates

Aligned reads spanning genomic positions of somatic mutations were re-genotyped using Samtools mpileup (v.1.9)^75^. Genotypes supported by ≥ 2 reads with a nucleotide quality score of ≥20 were reported, considering sites with two alleles as biallelic, those with three or four alleles as multiallelic. To simplify analysis and interpretation, multiallelic rates were only calculated for mutations from A or T nucleotides (C and G lesions contribute to both DEN1 and DEN2 signatures, have differing propensities for multiallelic variation, and the signatures vary in their contributions between tumours). The multiallelic rate is the fraction of mutations from A or T in a tumour that are identified as multiallelic in that tumour. Multiallelic rate was calculated (i) as an average for each tumour, and (ii) separately for each mutation asymmetry segment of each tumour.

### MRCA generation and post-MRCA selection

Following prior work, we estimated cell generation post-mutagenesis of the most recent common ancestor (MRCA) of the cells in the sequenced tumour based on the fraction of autosomal genomic segments that are multiallelic^14^. Genomic segments were defined per tumour using mutational asymmetry, as described above. After the first cell division post-damage, each segment will contain 2 lesion-containing strands, which will be diluted through subsequent divisions. The scenarios where one or both lesion strands of a segment are multiallelic are not readily distinguishable from the data, as in both cases the segment will appear as multiallelic. However, the fraction of non-multiallelic segments, where both lesion strands are lost, is directly observable from the data. We call a segment non-multiallelic if <4% of mutated sites are multiallelic. Let *p* be the fraction of autosomal lesion containing strands, and *q*=1-*p*. Assuming independent segregation of DNA copies per segment, the expected fraction of autosomal segments without multiallelism is *q^2^*, and thus we estimate *p* as 1-(fraction of segments without MA)^½^.

Tumours with >75% or 0% of segments showing multiallelism were assigned generation 1 and generation >=5 MRCAs, respectively. For the remaining tumours, we estimated the MRCA generation using maximum likelihood under the following rationale. At generation 1 post-mutagenesis, we expect 100% of segment copies to have a lesion containing strand and thus show multiallelism. At each subsequent generation, a segment loses one of its lesion-containing strands, and thus its ability to generate multiallelism with probability 0.5: hence the probability of retaining an individual lesion strand at generation *n* post mutagenesis is *2^(-n+1)^*. With independent segregation, the probability of no lesion strands in a segment at generation *n* is thus (1-*2^(-n+1)^*)^2^. In a tumour with *x* autosomal segments, assuming segment independence, the observed segment number without MA is thus distributed as Binomial(*x*, (1-*2^(-n+1)^*)^2^). Ranging over *n=2,3,4,5*, we selected the *n* value that maximises the likelihood of the observed data under this binomial model, resulting in an inference of the MRCA generation of the tumour.

To quantify post-MRCA selection we identified genomic segments where the two allelic copies of a chromosome have opposite mutational asymmetries, resulting in the segment as a whole being mutationally symmetric (−0.33 ≤ S ≤ 0.33). In these mutationally symmetric regions, T→N mutations relative to the reference genome forward strand represent mutations from one lesion strand and A→N, mutations represent the other lesion strand of the segment. Calculating multiallelic rates separately for T→N and A→N allows determination of the multiallelic rate resulting from each lesion strand of a segment. Under the rationale that unbalanced lineage contributions would result in higher variation in multiallelic rate (**Fig. 7a**), the coefficient of dispersion (the standard deviation divided by the mean) of the MA rates of these segments was taken as a metric of post-MRCA selection. Only lesion strands within mutationally symmetric regions and with MA rate ≥ 4% were considered. The MA coefficient of dispersion was calculated for n=397 mice, including tumours with inferred generation 1,2 or 3 MRCAs and that had at least 2 segments that satisfied the lesion strand filtering conditions. C3H tumours were classified as highly dispersed by using the non-C3H as a null-distribution, identifying tumours with MA coefficient of dispersion higher than the 95th percentile of MA coefficient of dispersion amongst non-C3H tumours.

### Genomic annotation

Genic annotation was obtained from Ensembl v.91^65^ for the corresponding C3H, BL6, CAST, and CAROLI reference genome assemblies (C3H_HeJ_v1, GRCm38, CAST_EiJ_v1 and CAROLI_EiJ_v1.1, respectively). Genomic repeat elements were annotated using RepeatMasker (v.20170127)^76^ with the default parameters and libraries for mouse annotation. Genomic coordinates were transformed between reference genomes using the HAL toolkit (v.2.3)^77^, utilising the UCSC mouseStrains_1509.hal multi-strain alignment with CAROLI_EiJ_v1.1 added using progressiveCactus (v.1.0.0)^78^. The functional consequences of mutations (e.g. amino acid change, splice site disruption, synonymous sequence change) were annotated using Ensembl Variant Effect Predictor (VEP, v.109.3) run in the ensemblorg/ensembl-vep docker container under Singularity; Ensemble v.91 annotation was used for the corresponding reference genome assembly. Phylogenetic distances between strains were calculated using four-fold degenerate (4D) sites, extracted from the HAL alignments using the HAL toolkit hal4dExtract function. 4D site rate calculations were performed using PHAST (v.1.3)^79^ phyloFit under the REV model.

### Driver discovery methods

Protein coding cancer driver genes were analysed using three computational methods: OncodriveFML (v.2.2.0)^29^ which identifies genes with a bias towards high impacting mutations, OncodriveCLUSTL (v.1.1.3)^30^ which identifies genes containing mutations that are linearly clustered in nucleotide sequence, and dNdScv (v0.1.0)^28^, which uses a maximum likelihood approach to quantify selection by identifying genes with an excess of nonsynonymous, essential splice site, or truncating mutations.

To increase the sample size, a single pan-strain analysis was performed using all DEN-treated DNs (n=581). Only consensus single nucleotide variants (SNVs) that mapped to all four genomes were kept (n=32,894,997). For the OncodriveFML and OncodriveCLUSTL analyses, mm10 was used as the reference genome. The dNdScv analysis was carried out four times, in each iteration mutations were projected onto the gene annotations of one of the four mouse strains.

To run OncodriveFML and OncodriveCLUSTL, protein coding sequences from all transcripts in protein-coding genes were merged together into their corresponding gene using pybedtools (v0.8.0)^80^.

OncodriveFML variant pathogenicity was assessed using GRCm38/mm10 SIFT scores (v.83)^81^. OncodriveFML was run on the pan-strain dataset using strain-specific coding DNA sequence (CDS) normalised trinucleotide background models. These mixed background models allow the method to account for strain-specific differences in the trinucleotide mutational probabilities. These models were computed using bgsignature (v0.2) (available at: https://pypi.org/project/bgsignature/). All other OncodriveFML parameters were kept as default. Genes with q-values < 0.1 were considered candidate drivers.

OncodriveCLUSTL was run using a strain-specific CDS normalised trinucleotide background model, as for OncodriveFML. In order to improve the accuracy of clustering signals detected in CDS regions, simulated mutations were randomly sampled within coding sequences (simulation mode “region_restricted”). Clusters were analysed by concatenating all subsequent CDS regions of a gene. All other parameters were kept as default. Genes with q-values < 0.01 bearing more than 15 mutations were considered candidate drivers.

To run dNdScv, genome references were built using the GTF (Ensembl v.91) annotations of the respective mouse strain. dNdScv was run with the parameters “cv=NULL” and “max_muts_per_gene_per_sample = 50”, and genes with q-values < 0.1 were identified.

### Driver event annotation and combinatorial analysis

Candidate driver events including point mutations, protein coding sequence disrupting indels, whole genome duplication and recurrent aneuploidies were encoded as binary present/absent variables for each tumour. For nucleotide substitution and indel mutations any “moderate” or “high” impact mutation annotation in an identified driver gene was considered to be a candidate driver mutation. Where VAF filtering of driver mutations was applied, a tumour was considered to have a driver mutation if the VAF of that mutation exceeded (c/2)/p where c is the cellularity estimate of the tumour and p is the expected ploidy of the driver locus (autosomal=2, X chromosome = 1, double those values in WGD tumours).

Evolutionary dependency (co-occurrence and mutual exclusivity) was scored using SELECT (v.1.6)^82^ with default parameters. The weighted mutual information (wMI) p-value was used for colour coding significance and false discovery rate (FDR) <0.1 used as multi-testing corrected threshold of significance^31^. Comparison of driver mutation proportions between strains were performed using the χ^2^ test implemented in the built-in chisq.test function of R, based on contingency tables of mutation counts.

### RNA-sequencing (RNA-seq)

Tumour RNA was isolated and purified as described above. RNA concentration was measured using a NanoDrop spectrophotometer (Thermo Fisher); RNA integrity was assessed on a Total RNA Nano Chip Bioanalyzer (Agilent).

Total RNA (1 μg) was used to generate sequencing libraries using the TruSeq Stranded Total RNA Library Prep Kit with Ribo-Zero Gold (Illumina 20020596 and 20020599), according to manufacturer’s instructions. Library fragment size was determined using a 2100 Bioanalyzer (Agilent). Libraries were quantified by qPCR (Kapa Biosystems). Pooled libraries were sequenced on a HiSeq4000 to produce ≥40 million paired-end 150 bp reads per library.

### RNA-seq data processing and analysis

Transcript abundances were quantified with Kallisto (v.0.43.1)^83^ (using the flag --bias) and a transcriptome index for each strain compiled from coding and non-coding cDNA sequences defined in Ensembl v.91^65^. Expression patterns among the four strains were identified through PCA by combining unnormalised counts for protein-coding genes with orthologs in GRCm38 (**Fig. 5a**), and per strain from all annotated protein-coding genes and long intergenic non-coding (linc) RNAs (**Fig. 5b**). The 500 most variable genes were selected in both cases. Transcripts per million (TPM) estimates were generated for each annotated transcript and summed across alternate transcripts of the same gene for gene-level analysis. Transcription start sites (TSS) for each gene were annotated with Ensembl (v.91) and based upon the most abundantly expressed transcript. RNA-seq data are available at ArrayExpress at EMBL-EBI under accession E-MTAB-8518^14^ and (accession pending).

### Differential expression analysis

Differential expression was called using DESeq2^84^ at the gene level after importing estimated counts per transcript from Kallisto using tximport^85^. Calling was restricted to protein-coding genes and lincRNAs. Pair-wise comparison of differentially expressed genes across strains was performed on the intersection of significantly differentially expressed genes from per-strain dysplastic nodule vs adult normal comparisons (p adjusted <=0.005) after mapping strain-specific IDs to GRCm38 (**Extended Data Fig. 7**).

### MAPK pathway gene expression analysis

Seventy genes in the MAPK pathway (KEGG pathway:mmu04010)^86^ were found to be differentially expressed in all strains in a *Braf*-driven tumour vs adult, age-matched normal liver tissue comparison. Fold changes of these genes, where significant, in *Hras* and *Egfr*-driven tumours was consistent with those in *Braf*-driven tumours (**Fig. 5c**). Differential expression was called as above, with restriction to tumours with identified driver genes. The expression of the 70 MAPK genes in the combined four-strains dataset described above were used in principal component analyses (**Fig. 5d**).

### Expected mutation rate modelling

To model the expected rate (mutations per million nucleotides) of a specific mutation in a focal gene (e.g. *Hras* Q61L is a T→A substitution in a GTT context with respect to the transcription template strand), we identified the 1,000 genes with the closest measured expression (transcripts per million) to the focal gene in the P15 liver RNA-seq. In aggregate for these nearest-neighbour genes, 192 category lesion-strand-resolved mutation rate vectors were calculated (see above) separately for transcription on (i) the lesion containing template and (ii) lesion containing non-template genic strands^17^. These rate vectors contain the rate per million nucleotides for each type of substitution (e.g. R_GTT→GAT_ for the template strand and R_AAC→ATC_ for the non-template strand lesions in *Hras* Q61L). For a set of tumours (e.g. n=370 C3H tumours), we counted the number of template (S_t_) and non-template (S_n_) retained lesion strands at a focal locus (e.g. *Hras* Q61) and used the ratio of template:non-template lesion strands to calculate an expected mutation rate for the specific focal mutation (e.g. expected rate μ = (R_GTT→GAT_S_t_+R_AAC→ATC_S_n_)/(S_t_+S_n_)). To estimate uncertainty we calculated the 95% confidence intervals from 100 bootstrap samples of the tumour set. The same bootstrap sampling of tumours provided 95% confidence intervals on the observed mutation counts.

The expected proportions of MAPK driver mutations for a set of tumours (e.g. n=370 C3H tumours) was calculated as μ_i_/∑μ_(1..n)_ where μ_i_ is the expected rate of the focal mutation and ∑μ_(1..n)_ the sum of the expected rates of for all considered MAPK driver mutations (*Hras* Q61R, *Hras* Q61L, *Hras* Q61K, *Kras* Q61R, *Egfr* F254I, *Braf* V637E). The 95% confidence intervals for the expected proportions were calculated from 100 bootstrap samples of the tumour set.

### Predicting the immune presentation of mutations

For every Variant Effect Predictor annotation of a substitution mutation (see above) that altered protein coding sequence, the corresponding amino acid change was edited into the translated peptide sequence and all possible overlapping 8, 9, 10 and 11 amino acid oligopeptides were presented to MHCflurry (v.2.1.1)^87^ and netMHCpan (v.4.1)^88^ for MHC class Iɑ affinity prediction. MHCflurry was used with pre-trained affinity models that correspond to the BL6 (H-2-Kb, H-2-Db) and C3H (H-2-Kk, H-2-Dk) class Iɑ MHC genes. No pre-trained MHCflurry models are available for CAST or CAROLI class Iɑ MHC genes.

The netMHCpan tool does allow for prediction even when the exact MHC class Iɑ gene was not included in the training set^88^. Strain specific MHC class Iɑ gene sequences for netMHCpan were defined as follows. The genomic sequences of the H2-D1 and H2-K1 genes, plus flanking sequence were queried from the mm39 genome reference using Bedtools: (v.2.30.0)^89^ (H2-D1 = chr17:35480951-35487230, H2-K1 = chr17:34213953-34220395). Orthologous sequence from each of the strains was found by querying the mm39 sequences against unannotated long-read based genome references (BL6 = C57BL_6NJ_v2, C3H = C3H_HeJ_v2, CAST = CAST_EiJ_v2, available from NCBI, BioProject = PRJEB47108) using BLAST (v.2.5.0)^90^. To predict the protein sequence, MHC class Iɑ protein sequences were obtained from UniProt^91^ (H-2-Db: P01899, H-2-Dk: P14426, H-2-Kb: P01901. H-2-Kk: P04223) and projected onto the H2-D1 and H2-K1 sequences using GeneWise^92^. Sequences were manually refined to ensure presence of start and stop codons, and resemblance to previously published exon boundaries. For the more divergent CAROLI strain, MHC class Iɑ protein sequences were obtained from NCBI RefSeq (BioProject: PRJNA387030); in contrast to the other strains, CAROLI has three annotated MHC class Iɑ genes (H2-K: XP_029329709.1, H2-D: XP_021009871.1, H2-L: XP_029326886.1).

The maximum affinity score (lowest rank_EL for MHCflurry; maximum pan_EL score for netMHCpan) of the multiple alternate overlapping peptides for each mutation was recorded and used in subsequent analysis. Note that mutations of every strain were scored against each of the MHC class Iɑ of every strain (e.g. BL6 mutations scored against each MHC class Iɑ gene from each of C3H, BL6, CAST and CAROLI). Affinity scores were quantile normalised for comparison to previously computed distributions of test peptides, as recommended^87,88^, and scores within the top 0.5% of test peptides were considered to be high affinity predicted binders. Normalised, maximum affinity scores were used for driver mutation immunogenicity prediction (**Extended Data Fig. 8**f).

As, for example, a C3H MHC molecule can only facilitate the immune mediated removal of mutations in C3H mice and not BL6, CAST or CAROLI mice, we tested for general evidence of immune editing by contrasting strain-matched versus non-strain-matched distributions of epitope affinity scores. We define the fraction of all (global) mutations that come from a focal strain as G_f_ (e.g. C3H mutations / (C3H + BL6 + CAST + CAROLI) mutations, for focal C3H). We produced a combined rank of affinity scores for global for each MHC class Iɑ gene, and take 1,000 mutation consecutive windows over the rank affinity scores, for each window calculating the (local) fraction of mutations from the focal strain L_f_. The relative enrichment (RE) of mutations from a strain in the window is calculated as RE=(L_f_-G_f_)/(L_f_+G_f_), a metric bounded (1,-1) where RE=0 for no enrichment. For the highest predicted affinity windows, immune editing would be predicted to lead to RE<0 (depleted mutations) where the focal strain matches the MHC class Iɑ gene used in affinity prediction. The results presented are restricted to mutations in expressed genes (median tumour expression TPM>1.0) as expression is expected to be a prerequisite for immune presentation, however the same conclusions can be drawn from analyses that do not filter on expression level. Confidence intervals (95%) were calculated from 10,000 random permutations of the affinity rank list. The analysis was repeated for all combinations of strain and each MHC class Iɑ gene, none showed compelling evidence supporting the extensive immune editing of neoepitopes; example analyses are shown (**Extended Data Fig. 8g**). Positive controls were provided by the computational subtraction of a defined percentage (4%, 2%, 1% or 0.2%) of high predicted affinity mutations, selected by sampling predicted weak and strong binders (top 5% normalised affinity) weighted by normalised affinity score (better predicted binders more likely to be selected for removal).

### Chromatin immunoprecipitation followed by high-throughput sequencing (ChIP-seq)

Livers from P15 mice were perfused *in situ* with PBS and then dissected, minced, cross-linked using 1% formaldehyde solution for 20 min, quenched for 10 min with 250 mM glycine, washed twice with ice-cold PBS, and tissue pellets were stored at –80°C. Tissues were homogenised using a dounce tissue grinder, washed twice with PBS, and lysed according to published protocols^93^. Chromatin was sonicated to an average fragment length of 300 bp using a Misonix tip sonicator 3000. To negate batch effects and allow multiple ChIP experiments to be performed using the same tissue, we pooled ten livers for each mouse strain; 0.5 g of washed dounced tissue was used for each immunoprecipitation. The following antibodies were used for each ChIP experiment: CTCF (rabbit polyclonal, Merck Millipore 07-729, lot 2517762, 20 μg); H3K4me3 (mouse monoclonal IgG clone CMA304, Merck Millipore 05-1339, lot 2780484, 10 μg); H3K27ac (rabbit polyclonal IgG, Abcam 4729, lot GR3187598, 10 μg). Immunoprecipitated DNA or input DNA (max 50 ng) was used for library preparation using the ThruPLEX DNA-Seq library preparation protocol (Rubicon Genomics, UK, R400676). Library fragment size was determined using a 2100 Bioanalyzer (Agilent). Libraries were quantified by qPCR (Kapa Biosystems). Pooled libraries were sequenced on a HiSeq4000 (Illumina) according to manufacturer’s instructions to produce paired-end 150 bp reads. All experiments were performed with a minimum of three biological replicates.

### ChIP-seq data processing and peak calling

To identify ChIP-seq positive regions, sequencing reads were trimmed to 50 bp and then aligned to their respective genome assemblies (BL6 = GRCm38, C3H = C3H_HeJ_v1, CAST = CAST CAST_EiJ_v1, CAROLI = CAROLI_EiJ_v1.1) using bwa (v0.7.17)^66^ using default parameters. Uniquely mapping reads from each library were selected for further analysis. Peaks were identified for each ChIP library using MACS (v.2.1.2)^94^ and matched input controls; for histone modifications, the “--broad” flag was used. For CTCF and histone modifications, all peaks with q-value <0.05 were included. We used the input libraries to filter spurious peaks associated with a high input signal using the GreyListChIP R package (v.1.36.0)^95^. Biologically-reproducible peaks were identified by merging ChIP-seq peaks (as defined above) from individual replicates and selecting those genomic regions found in two or more replicates.

Enhancers and promoters were defined using the sets of biologically-reproducible H3K4me3 and H3K27ac peaks following overlap rules defined previously^96^. Briefly, promoters were defined as H3K4me3 regions, with or without overlapping H3K27ac. Enhancers were defined as H3K27ac regions that did not overlap a H3K4me3 region. Finally, since regions of abnormal read coverage were masked for mutation detection (described previously), all ChIP-seq regions overlapping these regions were also removed from downstream analyses.

### Regulatory region mutation rate calculation

Genomic coordinates were transformed between reference genomes using the HAL toolkit (v.2.3)^77^, utilising the UCSC mouseStrains_1509.hal multi-strain alignment with CAROLI_EiJ_v1.1 added using progressiveCactus (v.1.0.0)^78^. Regions mapping to multiple scaffolds and overlapping alignments were removed, and the longest alignment for each region was identified. These 1:1 alignable regions were then overlapped with regulatory region annotation (above) in their corresponding strain.

For a given regulatory region type (CTCF binding sites, enhances, or promoters), we calculated the weighted-mean mutation rate of SNVs across all trinucleotide contexts. Mutation rate was calculated on a per-tumour basis as the fraction of each trinucleotide in the aggregated genomic span of a group of regions (e.g. promoters) that are mutated, weighted by the frequency of that trinucleotide in the genome (including regions masked for abnormal read coverage in that strain). To compare the influence of CTCF binding and regulatory site activity on mutation rate between strains, for each strain we calculated a relative enrichment metric between the bound/active sites and the shadow sites. Shadow sites are orthologous sequences to bound/active sites in other strains but not identified as bound/active in the focal strain (**Extended Data Fig. 6e**). Rates were calculated in aggregate over the tumours of a focal strains but individual binding and shadow sites were sampled with replacement 1,000 times to calculate bootstrap confidence intervals. The relative enrichment metric (*μRE*) was calculated as *μRE=(μ_active_-μ_shadow_)/(μ_active_+μ_shadow_); μRE>*0 shows active site mutation rate is greater than the shadow rate, *μRE<*0 shows the opposite, and *μRE=*0 denotes equal rates. Comparisons include separate calculations for (i) where the active sites are conserved across all four strains (4 way conserved) and (ii) where they are active in the focal strains but not detected as active in all four strains (partially conserved). Statistical tests for difference in mutation rate between active and shadow regions were implemented as Wilcoxon matched-pairs signed-rank test using the R wilcox.test function, with aggregate active and aggregate shadow regions within a tumour the matched-pairs; p-values were Bonferroni corrected for multiple testing (n=24 tests).

ChIP-seq data (FASTQ files and peak calls) are available in ArrayExpress at EMBL-EBI under accession E-MTAB-11959 and E-MTAB-14454.

### Assay for Transposase-Accessible Chromatin with high-throughput sequencing (ATAC-seq)

ATAC-seq protocol for the frozen tissue was used as described previously^97^, with minor modifications to the nuclear isolations steps. In Step 1: 1 ml of 1× homogeniser buffer was used instead of 2 ml, and in Step 4: douncing was performed with 30 strokes instead of 20)^17^. Pooled libraries were sequenced on a NovaSeq 6000 (Illumina) to produce paired-end 50 bp reads, according to the manufacturer’s instructions. Experiments were performed with at least 3 biological replicates.

ATAC-seq was also performed on DEN-treated BL6 tissues: 3 pairs of DEN-induced tumour and background non-tumour, 1 unpaired tumour, and 4 additional DEN-treated background liver tissue.

### ATAC-seq data processing and analysis

ATAC-seq data processing was performed using a custom-made Snakemake pipeline (v.6.1.1)^98^. Quality of raw reads was assessed using fastQC (v0.11.9)^99^, and adaptor sequences removed using cutadapt (v.2.6)^100^. Reads were aligned to their respective reference genome assemblies using bwa (v.0.7.17)^66^. Duplicate reads were marked using Picard (v.2.23.8)^101^.

Alignment filtering was performed with samtools (v.1.9)^75^ and reads overlapping genomic regions with abnormal read coverage (described previously) were removed. Reads aligning to mitochondrial DNA were excluded from further analysis. Read positions aligning to plus and minus strands were offset by +4bp and −5bp, as described previously^102^.

Peaks were called using macs2^94^ from all fragments, where ‘fragment’ refers to the inset between 5’ ends of read 1 and read 2, and sub-nucleosomal size fragments (<100bp). This was done for each sample separately and for a pool containing all replicates per condition.

Irreproducible Discovery Rate (IDR) method^103^ was used to determine a set of reproducible peaks. IDR was performed pairwise on replicates, using a subset of pooled peaks as reference. This subset contained pooled peaks that overlap at least 25% of the peak length in both replicates compared. Pooled replicate peaks that pass the IDR threshold of 0.05 in at least one pairwise comparison were deemed reproducible.

ATAC-seq data (FASTQ files and peak calls) are available in ArrayExpress at EMBL-EBI under accession E-MTAB-11780^17^ and E-MTAB-14144.

### Computational analysis environment

Quality control and alignment of WGS, ChIP-seq, ATAC-seq and RNA-seq data, and WGS variant calling was performed in a Linux cluster with LSF batch control. Except where otherwise noted, subsequent analysis was performed on a Linux cluster with Altair Grid Engine batch control, analysis in Conda environments and choreographed with Snakemake (v.7.32.4). Statistical analysis and data figure generation were performed in R^104^ (v.4.3.2).

### Data availability

Raw data files for all new datasets are available from Array Express, the European Nucleotide Archive (ENA), and BioImage Archive at the EMBL-EBI. WGS (BL6 and CAROLI) accession number from ENA: PRJEB15138. RNA-seq accession number from Array Express: pending. WSI (BL6 and CAROLI) accession from BioImage Archive: S-BSST384. ChIP-seq accession number from ArrayExpress: E-MTAB-14454. ATAC-seq accession number from ArrayExpress: E-MTAB-14144.

Previously published datasets were also analysed^14,17^: WGS (C3H and CAST) accession number from ENA: <u>PRJEB37808</u>. RNA-seq (C3H and CAST, P15 only) accession number from Array Express: <u>E-</u> <u>MTAB-8518</u>. WSI accession from BioImage Archive (C3H and CAST): <u>S-BSST383</u>. ChIP-seq (C3H CTCF) accession number from ArrayExpress: <u>E-MTAB-11959</u>. ATAC-seq (C3H) accession number from ArrayExpress: <u>E-MTAB-11780</u>.

### Key resources

The key reagents and resources required to replicate our study are listed in **Supplementary Table 2**. For externally sourced data, where applicable, URLs that we used can be found in the Git repository https://git.ecdf.ed.ac.uk/taylor-lab/lce-sd

## Acknowledgements and funding

We thank P. Bankhead for supervision of image processing; M. Roller and A. Yates for data curation; the CRUK Cambridge Institute Core facilities for their valuable contribution: CRUK Biological Resources (A. Mowbray), Preclinical Genome Editing (L. Young, S. Kupczak, M. Cronshaw, P. Mackin, Y. Cheng, and L. Hughes-Hallett), Genomics (F. Bowater and J. Hadfield), Bioinformatics (G. Brown, M. Eldridge, and R. Bowers), Histopathology and ISH (C. Brodie, L.-A. McDuffus, and J. Arnold), and Research Instrumentation (J. Gray); the EMBL-EBI Technical Services Cluster.

This work was supported by the Cancer Research UK Cambridge Institute core award (20412), MRC Toxicology Unit core funding programme grants (RG94521 and MC_PC_24012), MRC Human Genetics Unit core funding programme grants (MC_UU_00007/11, MC_UU_00007/16, MC_UU_00035/1, MC_UU_00035/2), and European Molecular Biology Laboratory core funding. IRB Barcelona is a recipient of a Severo Ochoa Centre of Excellence Award from Spanish Ministry of Science, Innovation and Universities (MICINN, Government of Spain) and is supported by CERCA (Generalitat de Catalunya). Support was also provided from specific research grants: Cancer Research UK strategic award (22398); Wellcome Trust (WT108749/Z/15/Z, WT202878/B/16/Z, 202878/Z/16/Z); PID2021-126568OB-I00 (CHEMOHEALTH) project, funded by the Spanish Ministry of Science (MCIN), AEI /10.13039/501100011033; ERC (615584, 788937); Helmholtz Society (DKFZ abteiling B270). This work made use of resources provided by the Edinburgh Compute and Data Facility (ECDF), including the IGC_Eddie3 high performance storage arrays (MC_PC-MR/X013677/1). Edinburgh Genomics is partly supported through core grants from NERC (R8/H10/56), MRC (MR/K001744/1) and BBSRC (BB/J004243/1). S.J.A. received a Wellcome Trust PhD Training Fellowship for Clinicians (WT106563/Z/14/Z and WT106563/Z/14/A), National Institute for Health Research (NIHR) Clinical Lectureship, and CRUK Clinician Scientist Fellowship (RCCCSF-May23/100001); J.C. was supported by a Wellcome Trust PhD Training Fellowship for Clinicians (WT223088/Z/21/Z) as part of the Edinburgh Clinical Academic Track (ECAT) programme. A.E. was supported by a UKRI Innovation Fellowship.

For the purpose of open access, the authors have applied a Creative Commons Attribution (CC BY) licence to any Author Accepted Manuscript version arising from this submission.

## Author contributions

SJA, MST and DTO coordinated the design and implementation of the project. SJA, FC, CF and DTO designed the experiments. SJA, FC and CF performed mouse experiments, tissue collection and processing. Genome sequencing was performed by SJA, CF and Edinburgh Genomics (JS-L). SJA and AMR performed RNA sequencing, SJA and MB conducted ChIP-seq and SC implemented ATAC-seq. SJA performed histological analysis. TFR and ML conducted sequence alignments and variant calling. SJA and MST analysed tumour susceptibility. JCV, CJA, OP, LT and MST analysed substitution and indel mutations. JL, JC and MST analysed copy number changes and whole genome duplication. CAP, CJA, VBK and MST performed driver mutation identification. JL and MST analysed driver gene frequency and immune editing. AE and SA performed gene expression analysis. VS performed regulatory site mutation rate analysis. JL, JH, MDN and MST designed and implemented measures of selection. MD, EK and IS performed supporting analyses. SJA, PF, MST and DTO obtained funding for the project. SJA, PF, NL-B, CAS, MST and DTO led the Liver Cancer Evolution Consortium. SJA, MST and DTO wrote the manuscript with contributions from JL, SA, JFH, MDN, VS, JCV, PF and CAS. All authors had the opportunity to edit the manuscript. All authors approved the final manuscript.

## Extended data

**Extended Data Fig. 1.**
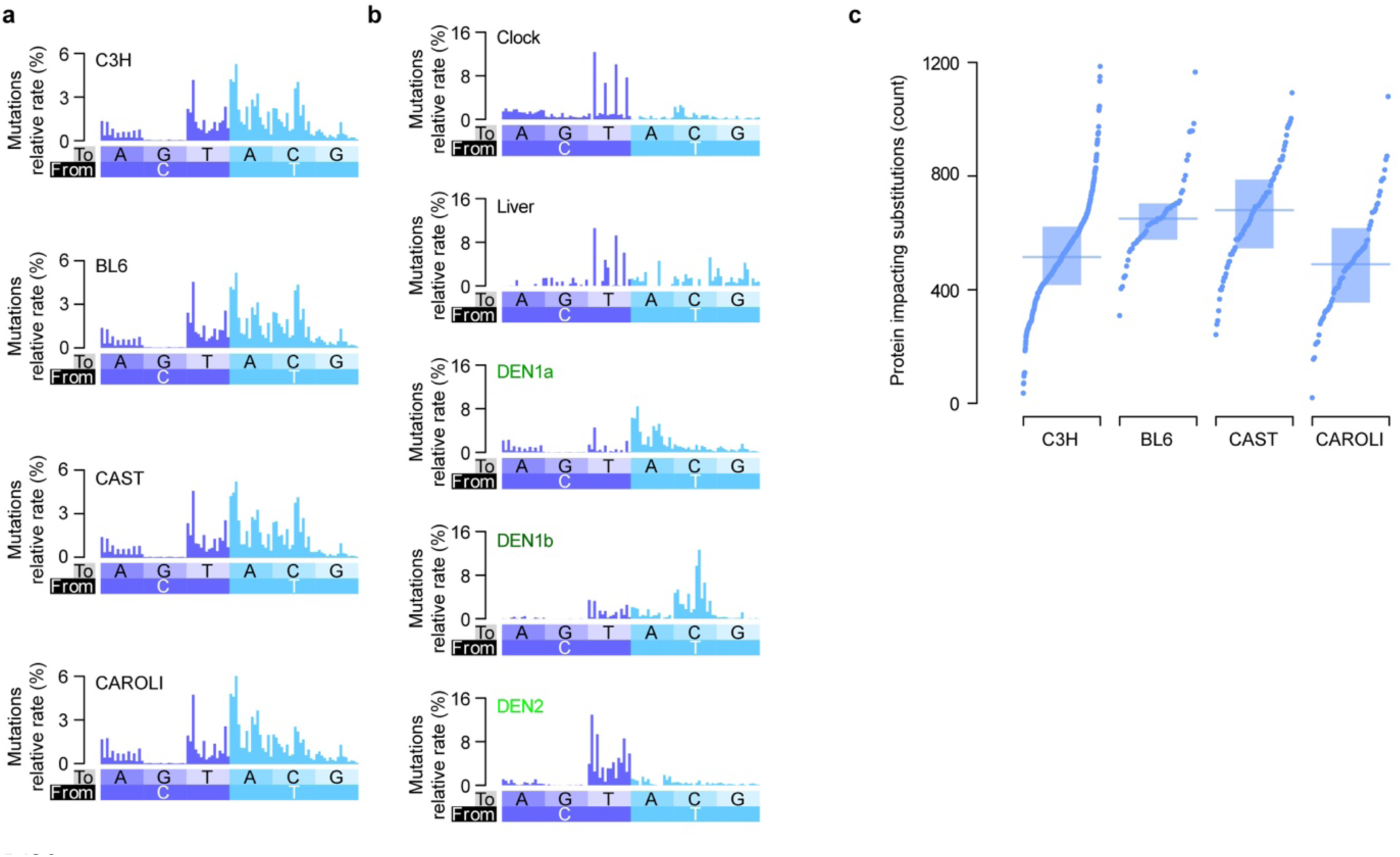
Single base substitution mutation signatures. **a,** Base substitution mutation rates calculated for DEN induced tumours for each strain. Substitution rates were calculated, stratifying by the substitution type and conditioned on the upstream and downstream nucleotide. Rates were calculated as the number of mutations divided by the number of trinucleotide occurrences, allowing direct comparison between genomes. Histograms were normalised so the total area of each histogram sums to 100%. Reverse complement substitutions were combined and are shown relative to substitutions from the pyrimidine nucleotide (e.g. ATG->ACG and CAT->CGT combined and shown as the former). **b,** Mutation signature deconvolution identifies five distinct signatures present amongst the 647 analysed genomes. The signature labelled Clock is predominant in non-DEN exposed samples (Fig. 2b) and closely matches expectations for a combination of the established human clock-like mutation signatures SBS1 and SBS5^60^. The signature labelled Liver is predominant in DEN-exposed, histologically normal samples which are also typified by low mutation rates (Fig. 2b). The remaining signatures are restricted to DEN exposed samples. Previously signatures DEN1 and DEN2 have been described for DEN induced C3H liver tumours^14^, here the DEN1 signature has split into DEN1a and DEN1b to capture the differences in the DEN induced mutation profile between CAROLI and the other strains, CAROLI demonstrating a reduction in T->C mutation (lower level of signature DEN1b). **c,** There are typically 400 to 800 protein impacting (non-synonymous, stop gain, stop loss or essential splice site) single nucleotide substitutions per DEN induced tumour. Horizontal lines show medians, shaded boxes the interquartile ranges.

**Extended Data Fig. 2.**
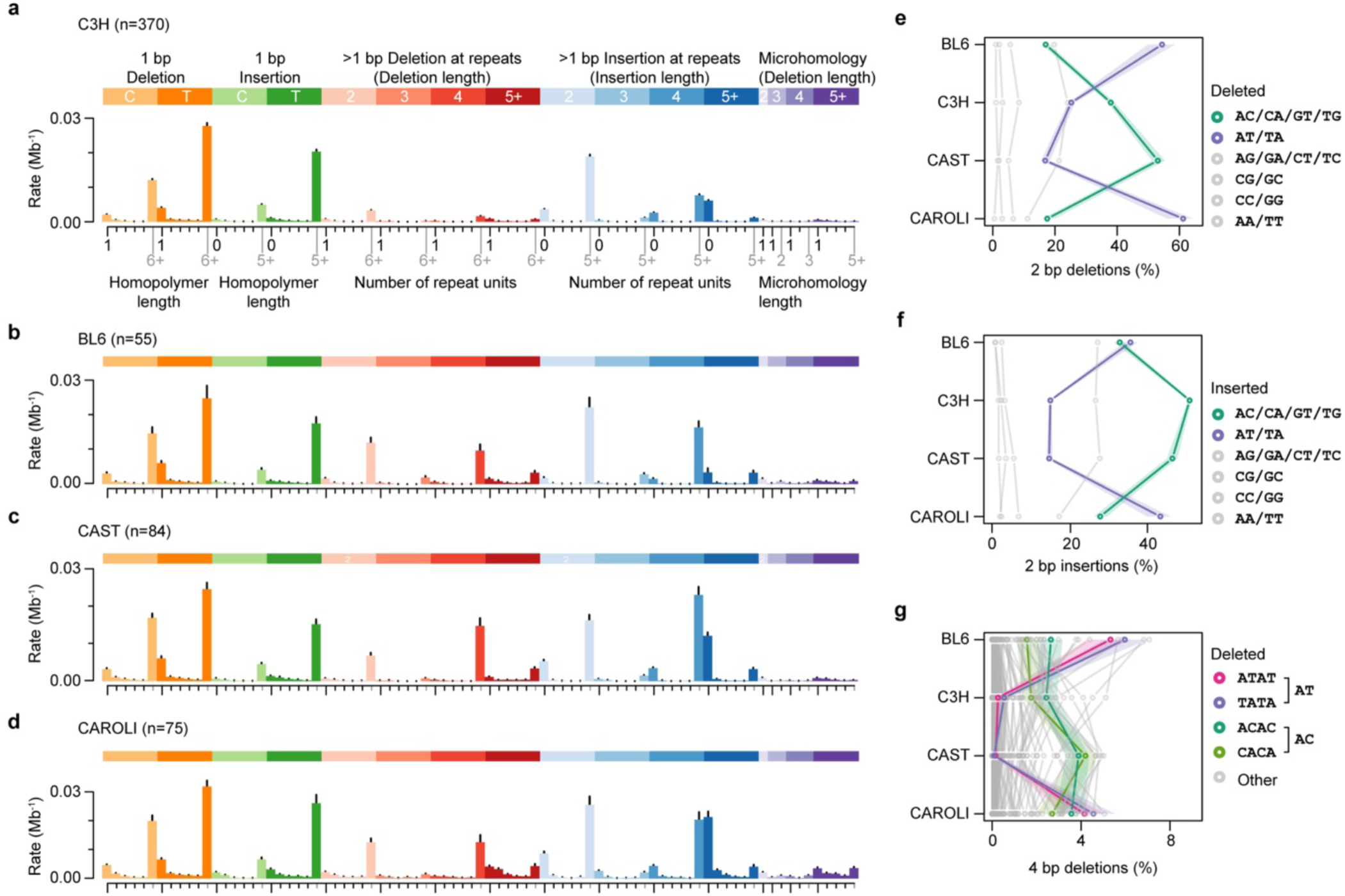
Indel mutation signature profiles of DEN-induced tumours. **a,** Insertion and deletion (indel) mutation profile for DEN induced C3H tumours. Mutations are shown as rates per Mb of genome sequence (y-axis), whiskers denote 95% confidence intervals from bootstrap sampling tumours. Eighty-three categories of indel mutation are defined (x-axis) stratifying by insertion/deletion, event length, and local sequence repetitiveness. The x-axis scale annotates the first (black) and last (grey) repeat measure for the colour coded groups of mutation category, intervening ticks are monotonically increasing. **b,** BL6 indel mutations plotted as **a**. **c,** CAST indel mutations plotted as **a**. **d,** CAROLI indel mutations plotted as **a**. **e,** The sequence composition of deleted dinucleotides differs between mouse strains, with AT (purple) the most deleted in BL6 and CAROLI and AC (green) in C3H and CAST; clustering does not reflect the phylogeny of the strains. For each strain (y-axis) the percentage of 2 bp deletions for the six possible dinucleotide categories (grouping circular permutations and reverse complements) is plotted (x-axis) with lines connecting corresponding points between strains. Shaded area around lines denotes the 95% confidence interval from bootstrap sampling of tumours. **f,** Dinucleotide insertions show similar strain biases to deletions. Plotted as **e**. **g,** strain biases for 4-nucleotide deletions show the bias for BL6 and CAROLI to delete AT dinucleotide containing sequences extends to larger deletions. Plotted as for **e-f** but considering all possible 4-nucleotide sequences.

**Extended Data Fig. 3.**
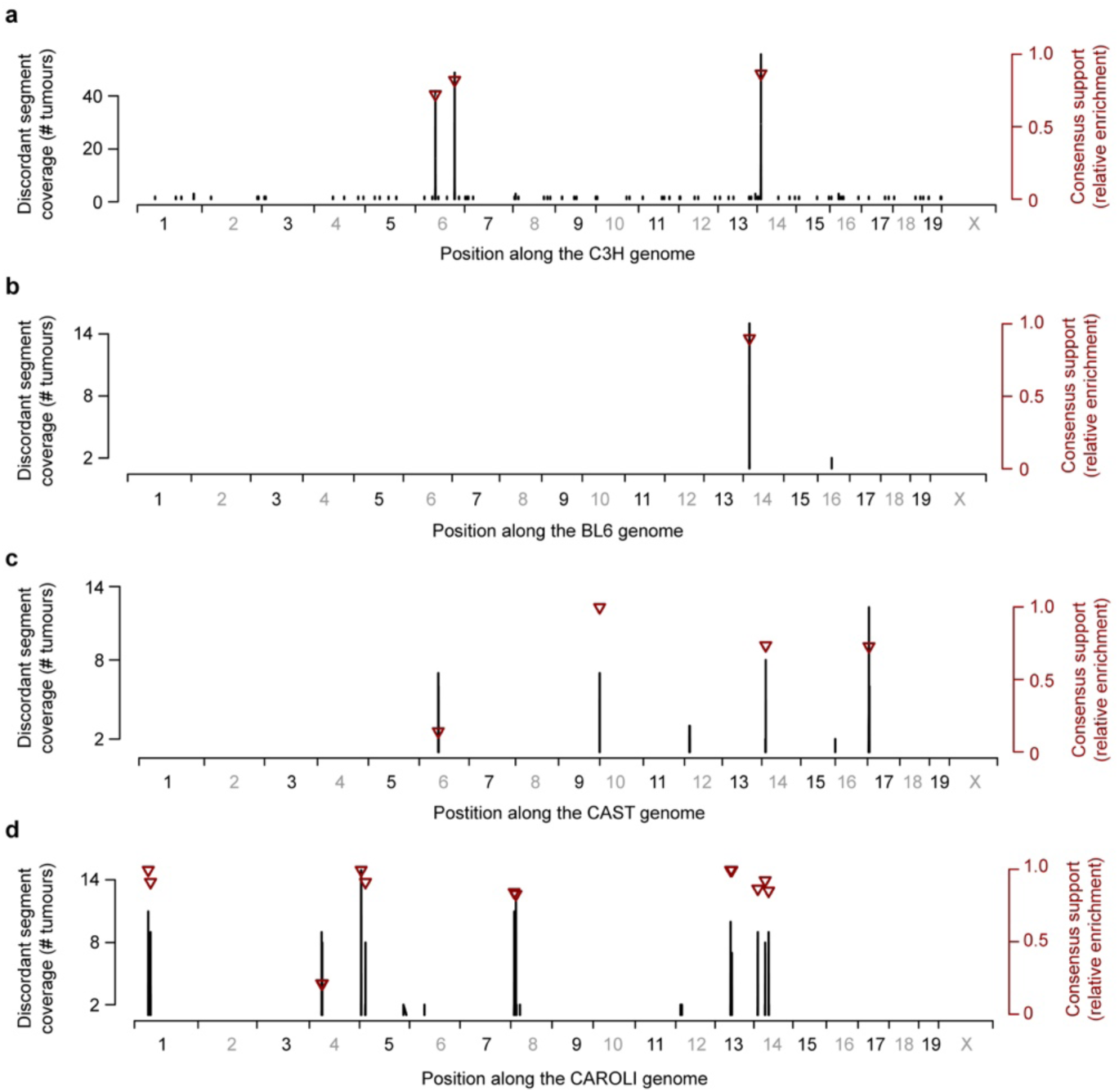
Sister chromatid exchange can be used to detect reference genome misassembly. **a,** Candidate C3H reference genome assembly errors (previously reported ^14^). Genome coordinates shown on the x-axis. Immediate switches between forward and reverse strand asymmetry are not expected on autosomes unless both copies of the chromosome have a sister chromatid exchange (SCE) event at equivalent sites. However, inversions and translocations between the sequenced genomes and the reference assembly are expected to produce immediate asymmetry switches. The discordant segment coverage count (black y-axis) shows the number of informative tumours (those with either forward or reverse strand asymmetry at the corresponding genome position) that suggest a tumour genome to reference genome discrepancy. Consensus support (brown y-axis) plotted as triangles shows the relative enrichment of informative tumours that support a genomic discrepancy at the indicated position (values > 0 indicate consensus support for misassembly). **b,** Candidate BL6 (GRCm38/mm10) reference genome assembly errors, plotted as per **a**. **c,** Candidate CAST reference genome assembly errors plotted as per **a**. **d,** Candidate CAROLI reference genome assembly errors plotted as per **a**. The only candidate error in the high quality BL6 reference genome (GRCm38/mm10), located on chromosome 14, corresponds to a proposed assembly error at the approximate orthologous position on chromosome 14 in each of the C3H, CAST, and CAROLI assemblies, suggesting the BL6 misassembly has propagated through the later-assembled genomes. The full list of assembly errors is provided in **Supplementary Table 4**.

**Extended Data Fig. 4.**
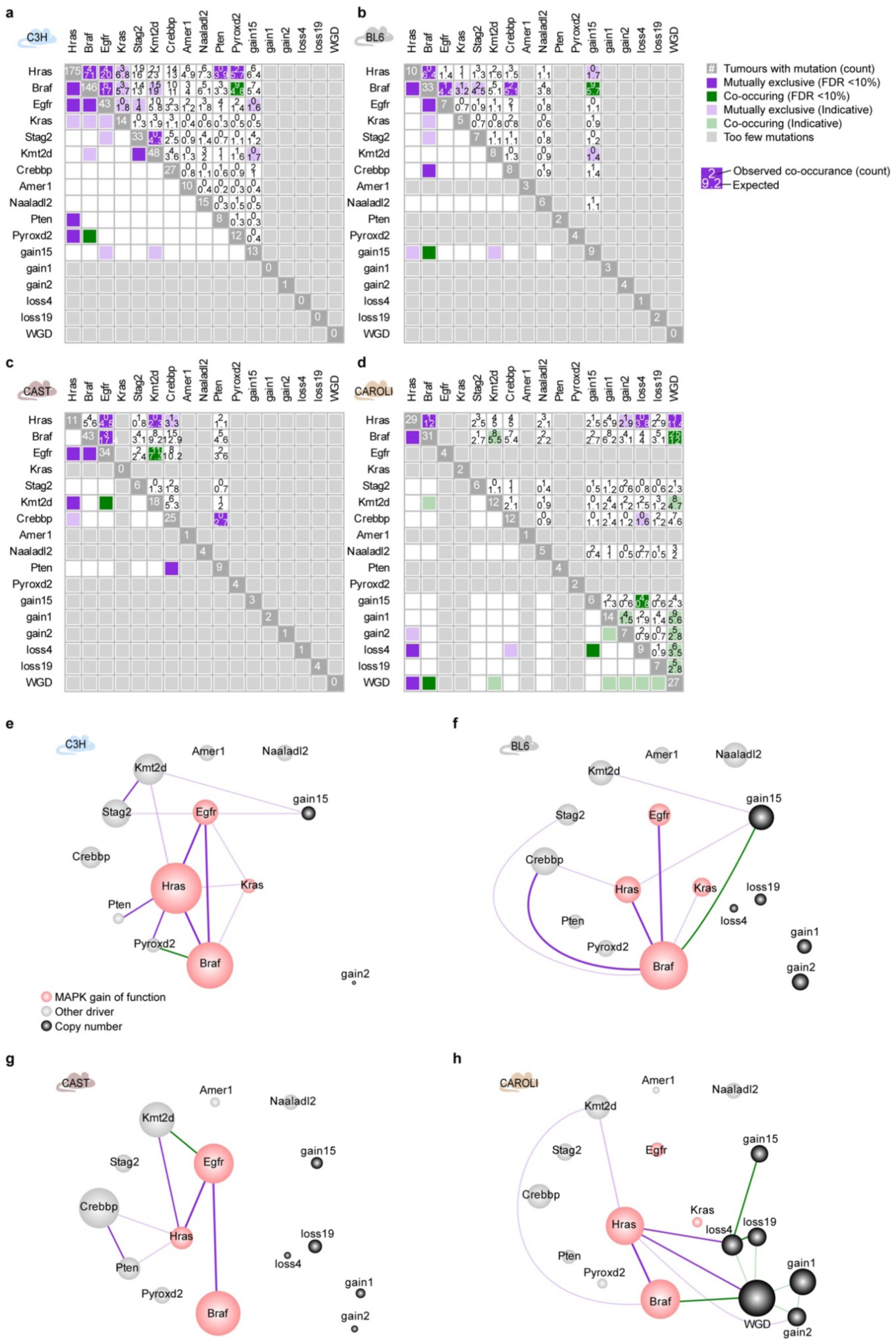
Strain comparison of driver mutation occurrence and interdependency. **a-d**, Co-occurrence and mutual exclusivity analysis for each strain: **a**, C3H; **b**, BL6; **c**, CAST; **d**, CAROLI. Dark green denotes significant (FDR<10%) co-occurrence of mutations within the same tumours, dark purple significant (FDR<10%) mutual exclusive occurrence. Statistical analysis by permutation^82^. Lighter shading denotes trend towards co-occurrence (green) and mutual exclusivity (purple). Some pairwise comparisons have insufficient power for statistical testing (light grey shading). Observed and expected co-occurrence counts are indicated in the upper triangle. **e-h**, Occurrence and interdependency graph for driver mutations in each strain. The area of each node is linearly scaled to the proportion of driver mutations, edges show co-occurrence and mutual exclusivity relationships identified and are coloured as in **a-d**. There is a high degree of consistency between strains in the driver mutations present and their co-occurrence/exclusivity relationships, but pronounced distortions in their relative frequencies.

**Extended Data Fig. 5.**
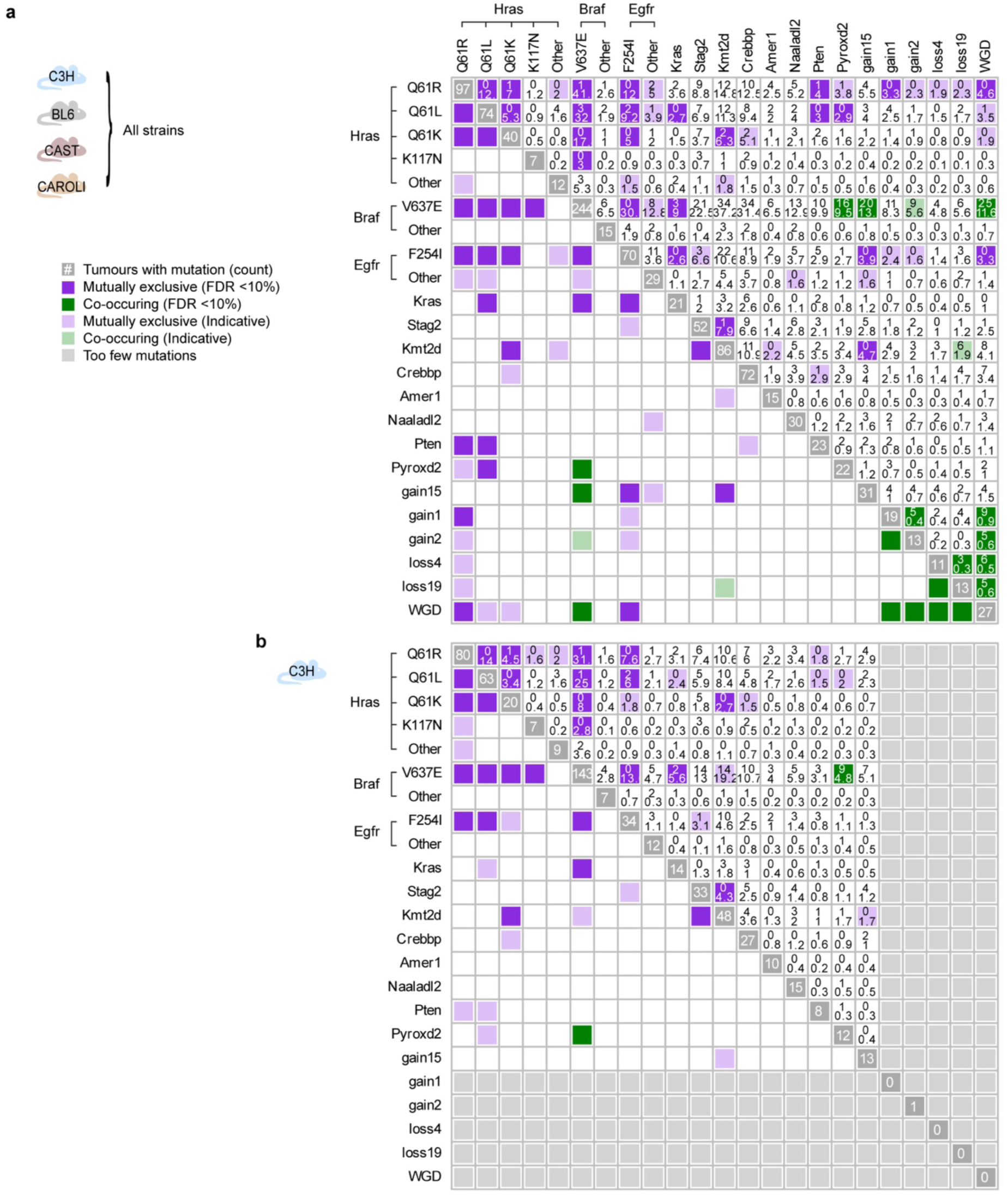
Driver mutation occurrence and interdependency extended to separately consider highly recurrent mutations. **a**, Aggregate analysis of mutations from all four strains where highly recurrent mutations in *Hras*, *Braf* and *Egfr* have been analysed separately, labelled by the specific amino acid changes. The label “Other” denotes other heterogeneous protein impacting changes in these genes; these show less strong co-occurrence and mutual exclusivity relationships indicating they may be mixed populations of both driver and non-driver mutations. **b**, As for **a**, but only considering mutations in C3H tumours.

**Extended Data Fig. 6.**
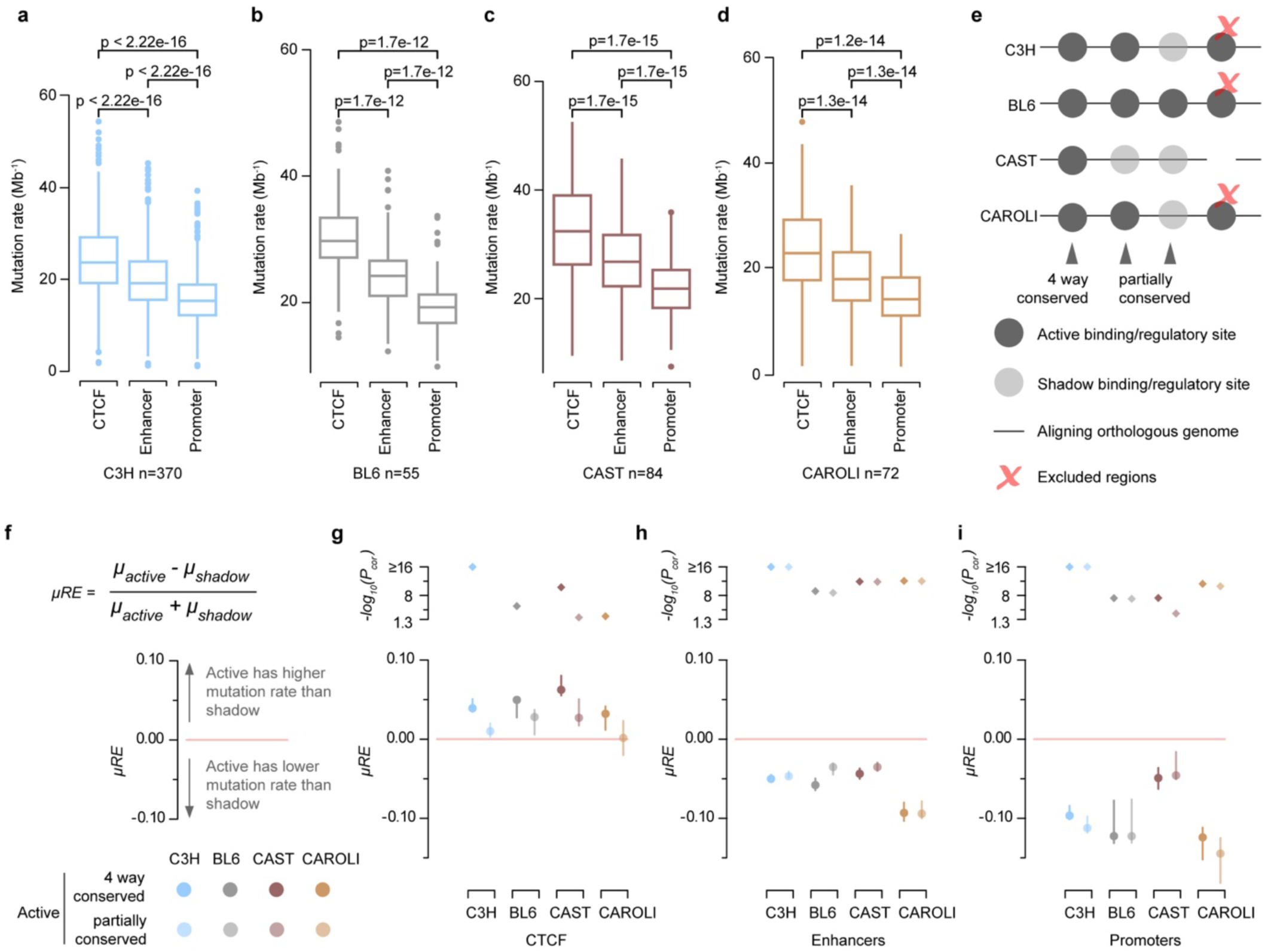
Mutation rates differ between regulatory site categories but show consistent behaviour between strains. **a**, Comparison of mutation rates between gene regulatory categories shows higher rates for CTCF binding sites than enhancers or promoters in C3H tumours. Mutation rates for ChIP-seq defined CTCF binding sites and histone modification defined active enhancers and promoters from P15 mouse liver. Rates calculated per tumour with outlier points indicated. The box defines the interquartile range and mid bar the median. Comparisons of mutation rate distribution between regulatory categories were calculated using a two-sided, paired Wilcoxon signed rank test. **b-d** Regulatory site mutation rate comparisons for BL6, CAST and CAROLI as for C3H in **a** shows a consistent rank of mutation rates with CTCF > enhancer > promoter. **e,** Using whole genome alignment, the cross strain conservation of regulatory activity was categorised. Orthologous regulatory sites active in all four strains were classed as 4 way conserved. Partially conserved sites were those where orthologous sequence could be identified in each strain but binding/histone modification was only identified in a subset of the four strains; the corresponding unbound regions are defined as shadow sites. Sites where orthologous sequence could not be identified in all strains were excluded from subsequent regulatory site analysis. **f,** The relative enrichment (RE) of mutation rates (μ) was calculated between active regulatory sites and shadow sites as defined in **e**. **g,** Active CTCF binding sites have higher mutation rates than shadow CTCF binding sites (μRE > 0), and 4 way conserved CTCF sites have higher mutation rates than partially conserved CTCF sites. Statistical significance (top panel) was calculated using two-sided, paired Wilcoxon signed rank test, between mutation rates for active and shadow binding sites, plotted p-values (y-axis) are Bonferroni corrected. Whiskers show bootstrap 95% confidence intervals. **h,** Active enhancers have lower mutation rates than shadow enhancers; calculated and plotted as in **g**. **i,** Active promoters have lower mutation rates than shadow promoters; calculated and plotted as in **g**.

**Extended Data Fig. 7.**
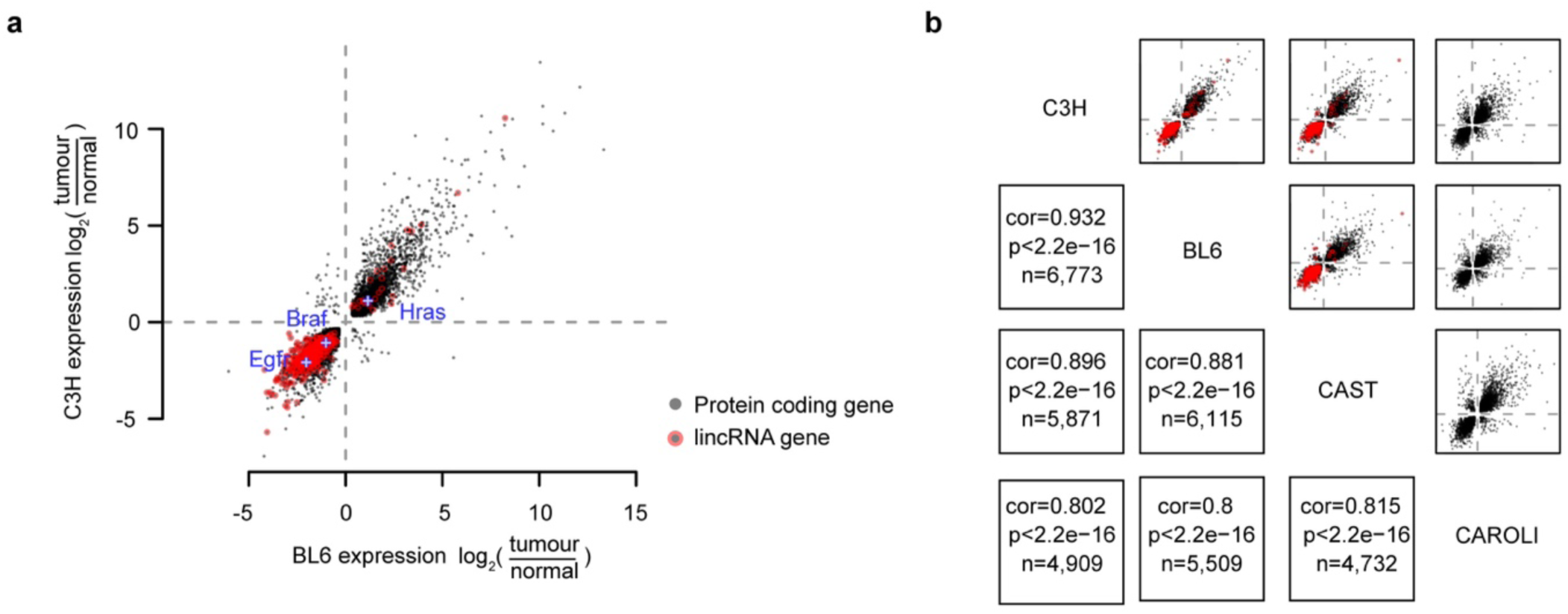
Consistent gene expression changes in tumour development. **a,** Changes in gene expression between age matched normal liver and liver tumours are highly correlated between evolutionarily divergent genomes. Each point denotes the expression change of a single orthologous gene pair between the BL6 and C3H genomes, only genes significantly differently expressed in both strains shown. In contrast to protein coding genes (grey), lincRNAs (red) are strongly biassed to reduced expression in tumours compared to normal liver. **b,** All versus all strain pairwise comparisons of differentially expressed genes; BL6 versus C3H is the same as (**a**). Pearson’s correlation coefficients, corresponding p-values, and counts of differential genes are shown. Equivalent lincRNA annotation is not available for CAROLI.

**Extended Data Fig. 8.**
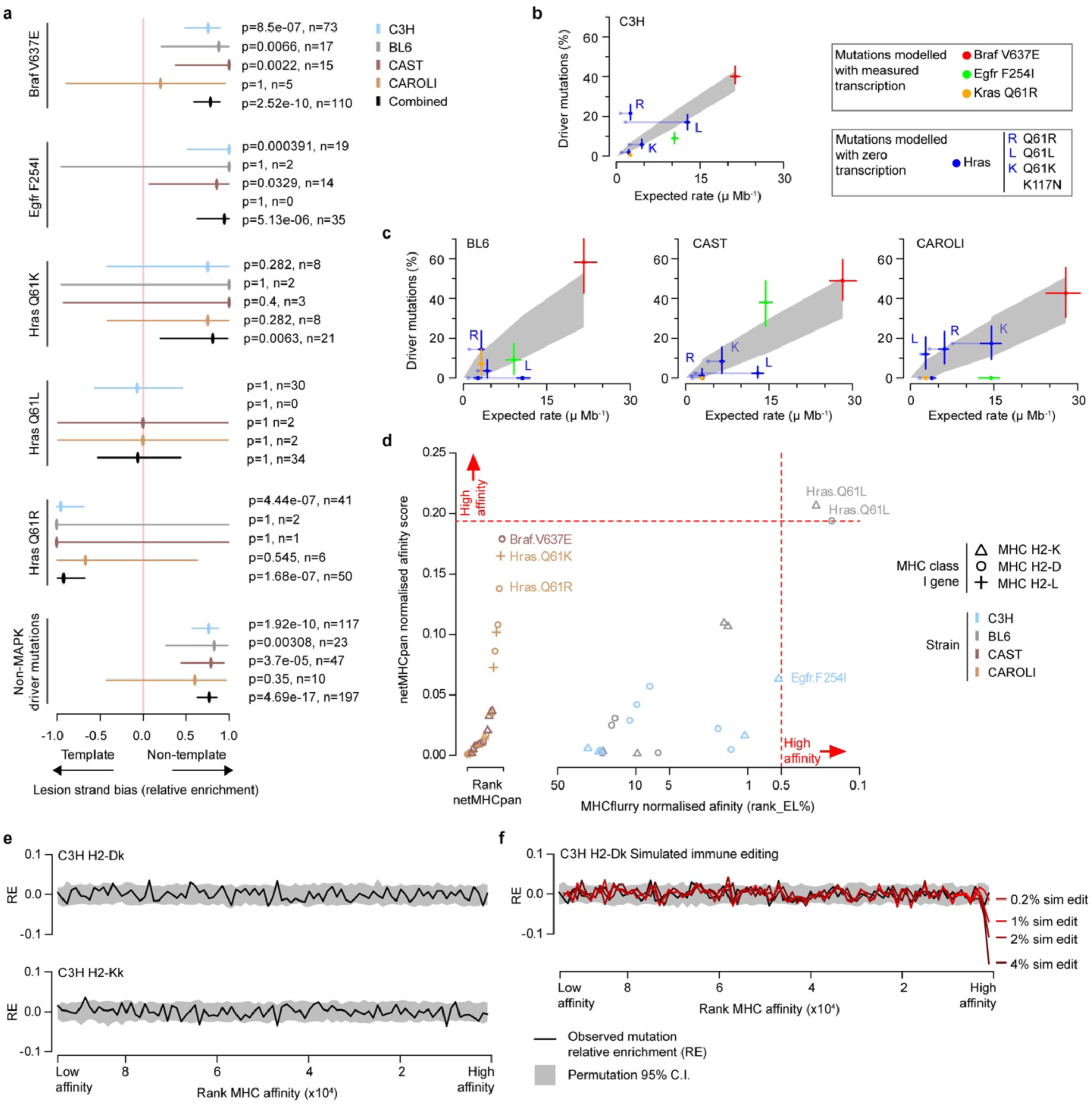
Mutational and immune influences on driver mutation bias. **a,** Bias of recurrent driver mutations to transcriptional template or non-template strand (x-axis) shown separately for each strain and the combined analysis aggregating across strains (black). Whiskers show the bootstrap 95% confidence interval. The number of tumours with each mutation (n) and nominal p-value (p; two-sided Fisher’s exact test) indicated. Lower panel shows aggregated analysis for mutations in non-MAPK pathway genes identified as containing driver mutations. **b,** Observed percentage of C3H tumours with each recurrently observed MAPK activating mutation (y-axis) compared with the expected mutation rate (μ, x-axis). Expected mutation rates were calculated based on lesion-strand resolved mutation rate and spectrum, adjusting for transcription template strand and gene expression level (Methods). Confidence intervals (95%, whiskers) for both observed driver mutation percentage and expected mutation rate were calculated from bootstrap sampling of tumours. Expected mutation rates for *Hras* were calculated assuming zero transcription (dark blue); the expected rates calculated using observed P15 bulk liver RNA-seq estimates of *Hras* expression shown (light blue; also shown in Fig. 6f) and solid line (light blue) connecting the alternative estimates. **c,** Observed versus expected driver mutation rates for BL6, CAST, and CAROLI, respectively, shown as for **b**. **d,** Predicted maximum affinity of recurrent driver mutation containing peptides for MHC class I molecules. Each of the mouse strains has two (C3H, BL6, CAST) or three (CAROLI) class I genes. For netMHCpan (y-axis) each mutation was scored for each MHC class I gene in each strain, whereas MHCflurry could only be applied to BL6 and C3H genes for which trained models exist. Scores for both netMHCpan and MHCflurry were quantile normalised as recommended and thresholds for high affinity binding indicated (right and above the red dashed lines). Only *Hras* Q61L in BL6 is predicted to be a high affinity binding peptide. **e,** For a defined MHC class I gene from an indicated strain (top, H2-Dk in C3H), all protein altering base substitution mutations from all four mouse strains were scored for their maximum binding affinity (MHCflurry). Binding scores were ranked and consecutive windows of 1,000 ranked scores were tested for the relative enrichment/depletion (RE) of strain-matched mutations compared to random expectation (C3H mutations when considering a C3H MHC gene). Immune editing of high affinity peptides is expected to show a depletion of strain-matched mutations (RE<0) at high affinity, compared to non-strain-matched mutations. Confidence intervals (95%, grey area) were calculated from permuted affinity ranks. Example analyses from C3H shown, equivalent analysis for each MHC locus in each strain was performed but no compelling evidence of substantial immune editing was evident. **f,** Immune editing simulated by computational subtraction of high MHC affinity strain-matched mutations from the observed distribution of C3H MHC gene H2-Dk affinity scores, simulating from 0.2% (bright red) to 4% (dark red) of protein altering mutations being removed probabilistically dependent on affinity score. This indicates that in the bulk analysis of mutations we have power to detect ≥1% of protein altering mutations removed. However, BL6 *Hras* Q61L is predicted to be in the top 0.2% of affinity scores, which is not adequately powered in this dataset (bright red).

**Extended Data Fig. 9.**
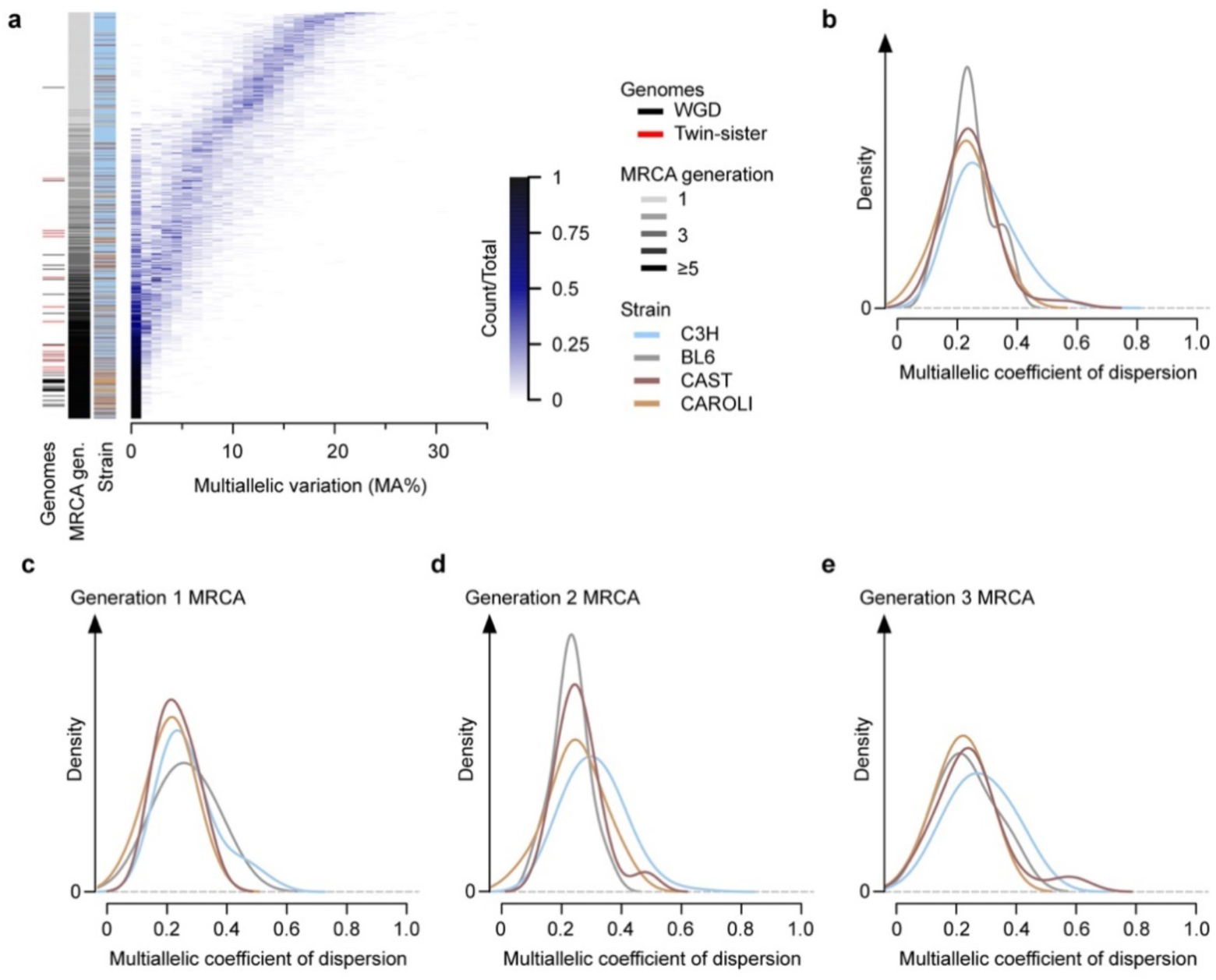
Subclonal dynamics of tumour development. **a,** Heatmap summary of multiallelic (MA) rate distribution for genomic segments in each tumour (rows), ranked by mean MA rate plotted in consecutive 1% bins (x-axis). Annotation of MRCA generation post-mutagenesis is based on the fraction of MA segments. Whole genome duplicated (WGD, black) and cases where both daughters of the originally mutagenised cell contributed to the sequenced tumour (twin-sister, red) are indicated. **b,** Density distribution analysis suggests a subpopulation of C3H tumours (19% estimated by comparison with the 95% quantile of non-C3H tumour dispersion) have higher multiallelic coefficients of dispersion than found for other strains. **c-e,** Stratifying analysis by the MRCA generation shows that all strains show a similar distribution of multiallelic coefficient of dispersion, confirming that the overall analysis (**b**) is not systematically confounded by strain differences in MRCA generation. A consistent trend is observed for C3H to be skewed to a higher dispersion than most other strains for each MRCA generation.

**Supplementary Table 1** | Table of tumours sequenced containing key metadata (Excel file).

**Supplementary Table 2** | Table of key resources, software, and datasets (Excel file).

**Supplementary Table 3** | List of DEN-naive mice assessed for the presence of tumours (Excel file).

**Supplementary Table 4** | List of reference genome mis-assemblies (Excel file).sample

